# The HORMA-domain protein ASY1 recruits the plant-specific cyclin SDS to promote DMC1-mediated meiotic recombination

**DOI:** 10.1101/2025.05.13.653901

**Authors:** Bowei Cai, Xinjie Yuan, Yanru Luo, Fei Cao, Yashi Zhang, Jixin Zhuang, Miaowei Geng, Lei Chu, Arp Schnittger, Chao Yang

**Affiliations:** National Key Laboratory of Crop Genetic Improvement, Hubei Hongshan Laboratory, Huazhong Agricultural University, Wuhan, 430070, China; Department of Developmental Biology, Institute of Plant Science and Microbiology, University of Hamburg, Hamburg, 22609, Germany

## Abstract

The chromosome axis is a proteinaceous scaffold essential for crossover formation in meiosis. While the axis associated HORMA-domain protein ASY1 is known to be necessary for the localization of the recombinase DMC1 to the axis, the molecular mechanism linking ASY1 to DMC1 loading remains unclear. Here, we demonstrate that ASY1 directly recruits the plant-specific cyclin SDS to chromosomes, which ensures efficient DMC1 localization and interhomolog recombination. Live-cell imaging and immunodetection revealed that SDS localizes to chromosomes from leptotene to zygotene and dissociates from chromosomes after pachytene resembling DMC1 dynamics. Biochemical assays demonstrated a direct interaction between ASY1 and SDS. Further analyses revealed that ASY1’s closure motif, a conformational regulatory element, is required for SDS recruitment despite being dispensable for ASY1’s own chromosome association. Moreover, unlike in *sds* mutants, depletion of the DMC1 inhibitor FIGL1-FLIP complex failed to restore DMC1 foci in *asy1* mutants, positioning ASY1 upstream of both positive (SDS) and negative (FIGL1-FLIP) regulators of DMC1. These findings establish ASY1 as a key coordinator of meiotic recombination, bridging chromosome axis architecture with recombinase dynamics to control crossover formation and genome stability.

## Introduction

Crossover formation, resulting from homologous recombination in meiosis, is a fundamental process for producing genetically different gametes in sexually reproducing organisms and is required for correct segregation of homologous chromosomes during meiosis (Martini et al. 2006; Zickler and Kleckner 2023).

The success of meiotic recombination relies on a series of tightly regulated steps, including the formation of programmed double-strand breaks (DSBs) and their subsequent processing, homology search, strand invasion, and the resolution of recombination intermediates (Zickler and Kleckner 2015; Lange et al. 2016). These steps are catalysed by a suite of enzymes with the support of cofactors, which function within the context of a chromatin-organizing structure known as the chromosome axis (Carballo et al. 2008; West et al. 2019; Yang et al. 2022).

The chromosome axis is a proteinaceous structure that assembles along the entire length of the sister chromatids between homologous chromosomes during early meiosis, serving as a basic scaffold for key meiotic events during prophase I, particularly homologous recombination (Zickler and Kleckner 1999, 2015; Blat et al. 2002; Panizza et al. 2011; Hunter 2015; Wang and Copenhaver 2018; Ito and Shinohara 2023; Chu et al. 2024). The axis organizes sister chromatids into a structure of linear DNA loop-arrays, facilitating efficient DSB formation, homology search and interhomolog-biased DSB repair, and synaptonemal complex assembly (Hollingsworth and Ponte 1997; Niu et al. 2005; Carballo et al. 2008; Goodyer et al. 2008; Hunter 2015; Xue et al. 2019; Lambing et al. 2020a, 2020b).

Key components of the chromosome axis have been identified that include (1) the cohesin complexes that encircle sister chromatids and are thought to provide a structural foundation for axis assembly across species including plants (Bhatt et al. 1999; Golubovskaya et al. 2006; Shao et al. 2011); (2) the HORMA domain-containing proteins (HORMADs), such as ASYNAPSIS1 (ASY1) in Arabidopsis (homolog of Hop1 in yeast and HORMAD1/2 in mammals); (3) coiled-coil proteins known as the ‘axis core’, e.g., ASYNAPSIS3 (ASY3) (homolog of Red1 in yeast and SYCP2 in mammals) and ASYNAPSIS4 (ASY4) (homolog of SYCP3 in mammals), which are required for proper ASY1 localization and axis integrity (Hollingsworth and Johnson 1993; Lammers et al. 1994; Ross et al. 1997; Armstrong et al. 2002; Wojtasz et al. 2009; Fukuda et al. 2010; Ferdous et al. 2012; Chambon et al. 2018). According to studies from many species, such as yeast, mammals, worms, and plants, the HORMAD proteins that localize on top layer of the axis, play a crucial role in meiotic recombination, especially during DSB formation, homology search, and strand invasion, thus also contributing to the homologous pairing and synapsis (Börner et al. 2004; Wojtasz et al. 2009; Shin et al. 2010; Daniel et al. 2011; Mercier et al. 2015; Lambing et al. 2017). For example, Arabidopsis *asy1* mutants exhibit severe defects in homology search, pairing, and bivalent formation, which is likely due to impaired chromosome association of the recombinase DMC1 (Sanchez-Moran et al. 2007; Lambing et al. 2020a; Pochon et al. 2023).

DMC1 is expressed specifically during early meiosis and is the main recombinase that promotes the homology search and strand invasion during meiotic DSB repair, ensuring crossover formation (Kurzbauer et al. 2012; Da Ines et al. 2013; Wang et al. 2016). DMC1, along with the recombinase RAD51 that functions in both mitotic and meiotic DSB repair, exerts its role by binding to 3’ single-strand DNA (ssDNA) overhangs to form nucleoprotein filaments. These filaments search for a matching sequence on the homologous chromosome, facilitating strand invasion, exchange, and homologous pairing (Brown and Bishop 2015; Hinch et al. 2020). However, the molecular mechanisms by which ASY1 regulates DMC1 localization remain poorly understood.

The loading of DMC1 onto chromosomes depends on factors that mediate DSB formation, such as SPO11 and MTOPVIB (Romanienko and Camerini-Otero 2000; Vrielynck et al. 2016). However, inactivation of *ASY1* in Arabidopsis does not, at least not obviously, affect DSB formation (Sanchez-Moran et al. 2007; Lambing et al. 2020a; Pochon et al. 2023), demonstrating that ASY1 likely does not affect DMC1 localization through regulation of DSB formation. In addition to proteins directly or indirectly involved in DSB formation, other factors that positively and negatively regulate the chromosome association of DMC1 in plants have been identified.

The positive DMC1 regulators include BRCA2 (Breast Cancer Susceptibility Protein 2), RAD51 and its paralogs, as well as SDS (SOLO DANCERS) (Bleuyard and White 2004; De Muyt et al. 2009; Tang et al. 2014; Zhang et al. 2015; Martinez et al. 2016). BRCA2 facilitates the formation of DMC1/RAD51 nucleoprotein filaments on ssDNA overhangs (Liu et al. 2010; Seeliger et al. 2012; Martinez et al. 2016). Arabidopsis *brca2* (*brca2a brca2b*) mutants are sterile and exhibit defective meiosis with neither DMC1 nor RAD51 foci being present anymore, resulting in a failure of DSB repair and a lack of recombination (Seeliger et al. 2012; Martinez et al. 2016). RAD51 and its paralogs, such as RAD51C and XRCC3, play a role in stabilizing the DMC1-ssDNA filaments and in their absence, DMC1 fails to localize at DSB sites (Tang et al. 2014; Zhang et al. 2015; Su et al. 2017; Jing et al. 2019). Additionally, the plant-specific cyclin SDS has been identified as a crucial promoter of DMC1 localization (De Muyt et al. 2009). In Arabidopsis *sds* mutants, DMC1 is not localized to the chromosomes, producing a phenotype similar to that of *dmc1* mutants with 10 univalents present in metaphase I meiocytes (Azumi et al. 2002; De Muyt et al. 2009). While both BRCA2 and RAD51 interact directly with DMC1(Siaud et al. 2004; Zhang et al. 2023), the molecular mechanism by which SDS regulates DMC1 loading onto meiotic chromosomes remains poorly understood.

Two complexes act as negative regulators of DMC1 localization, modulating its activity to avoid erroneous recombination events. First, the SMC5/6 complex, which shares features with cohesin and condensin, is recruited onto DNA to promote replication and DSB repair in both mitotic and meiotic cells. This complex inhibits excessive DMC1 loading onto meiotic chromosomes by sequestering DMC1 in the nucleoplasm via direct binding (Chen et al. 2021). Mutations in subunits of the SMC5/6 complex, e.g., ASAP1 and SNI1 in Arabidopsis (the homologs of Nse5 and Nse6 in yeast, respectively), lead to an increase in DMC1 foci (Chen et al. 2021).

Second, the AAA-ATPase FIGL1 (FIDGETIN-LIKE 1) and its interacting partner, FLIP (FIDGETIN-LIKE-1 INTERACTING PROTEIN) form a complex that interacts directly with RAD51 and DMC1, inhibiting their localization on meiotic chromosomes. This repression of DMC1 localization prevents the stand invasion step of homologous recombination during DSB repair (Girard et al. 2015; Fernandes et al. 2018). The *figl1* and *flip* mutants in Arabidopsis exhibit a hyper-recombination phenotype during meiosis, characterized by an elevated number of RAD51 and DMC1 foci and an increased frequency of crossovers. Notably, the FIGL1-FLIP complex antagonizes two DMC1 positive regulators, SDS and BRCA2. Mutations in *FIGL1* or *FLIP* can partially restore DMC1 focus formation, synapsis, and crossover formation in *sds* and *brca2* mutants (Girard et al. 2015; Fernandes et al. 2018; Kumar et al. 2019).

In this study, we aim to investigate how the axis protein ASY1 facilitates the chromosome localization of DMC1 and its relationship with both positive and negative regulators of DMC1. We characterized the expression and localization of SDS during meiosis and found that ASY1 recruits the positive regulator SDS through direct binding, which ensures the accurate chromosome localization of DMC1, thereby promoting inter-homologous DSB repair and crossover formation. Additionally, we observed that depletion of FIGL1 and FLIP does not, even partially, rescue the defect in DMC1 focus formation in *asy1* mutants, suggesting that ASY1 acts upstream of the FIGL1-FLIP complex in DMC1 loading onto meiotic chromosomes. Therefore, our study reveals a mechanism by which ASY1 promotes inter-homologous recombination through SDS recruitment to ensure proper DMC1 loading on chromosomes.

## Result

### ASY1 facilitates the chromosome loading of DMC1

Previous studies using immunostaining have shown that DMC1 localization on chromosomes is strongly compromised in the absence of ASY1 (Sanchez-Moran et al. 2007). We confirmed this finding in our immunostaining analysis of DMC1 in male meiocytes of *asy1* mutants (Figure 1A and B). This suggests that ASY1 is required for either the efficient chromosome loading or the protein stability of DMC1, or both.

**Figure 1.**
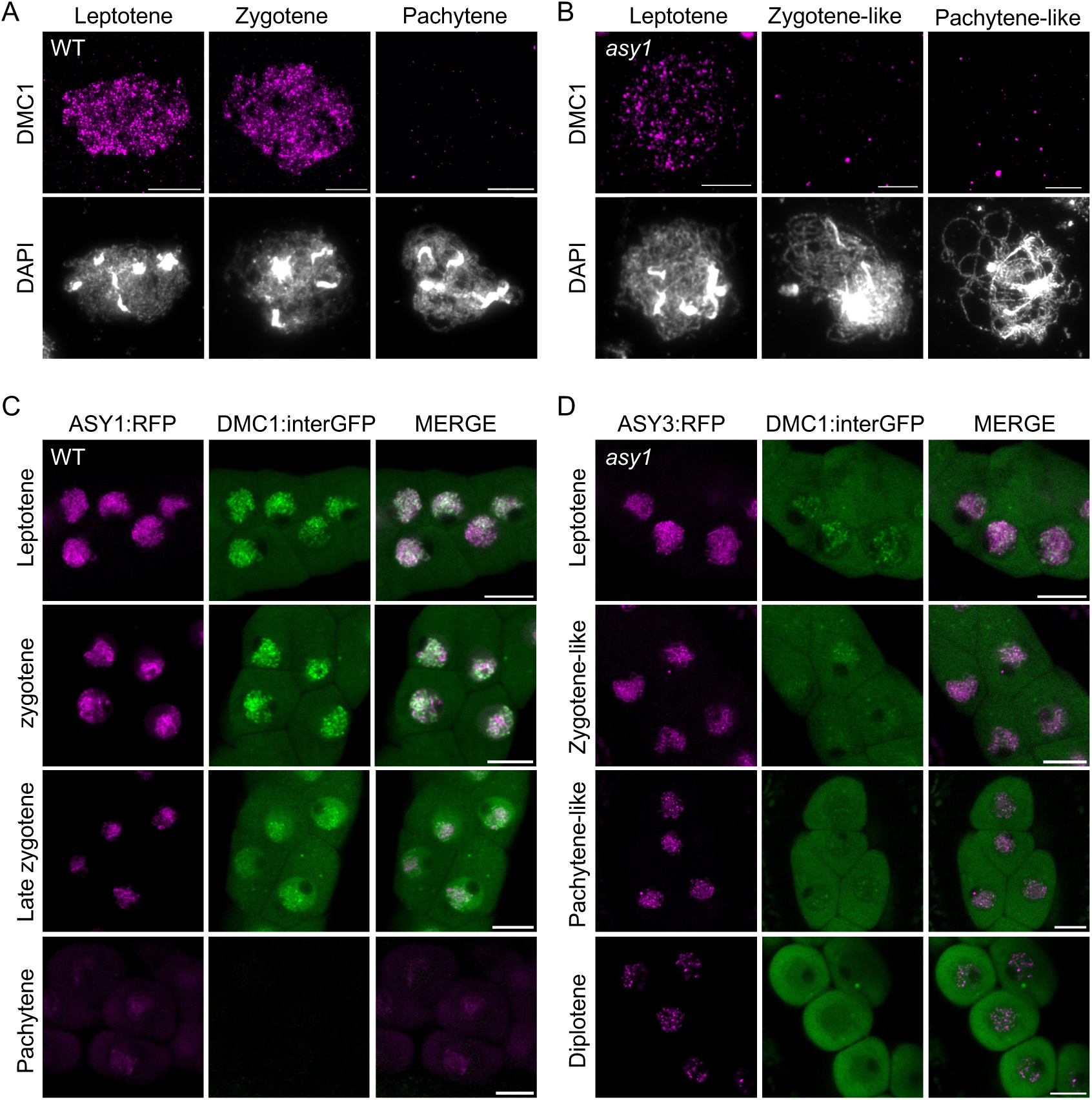
Chromosome localization of DMC1 is compromised in Arabidopsis *asy1* mutants. A-B: Immunostaining of DMC1 on spread chromosomes from male meiocytes of the wild-type (WT) (A) and *asy1* mutant (B) plants. Scale bars=10 µm. C-D: Live-cell confocal images showing the expression and localization patterns of DMC1:interGFP together with ASY1:RFP or ASY3:RFP in male meiocytes of wild-type (C) and *asy1* mutant (D) plants. Scale bars=10 µm.

To address this question, we sought to examine DMC1 localization in live meiocytes of *asy1* mutants using a fluorescent reporter and a previously established live-cell imaging set up (Prusicki et al. 2019). According to previous studies, both the N- and C-terminally GFP-tagged DMC1 reporters are not functional (Da Ines et al. 2022). Therefore, we generated a new reporter by inserting a GFP tag internally at the non-conserved region between the amino acids of 5 and 6 of DMC1 (*PRO_DMC1_:DMC1:interGFP*, hereafter *DMC1:interGFP*). Importantly, we found that this construct could rescue, albeit not fully, the sterility of *dmc1* mutants (seed setting: 18.40 ± 1.50 in *DMC1:interGFP* (*dmc1*) vs 1.00 ± 1.18 in *dmc1*, and 62.67 ± 2.45 in wildtype), suggesting that this *DMC1:interGFP* reporter retains substantial functionality (Supplemental figure 1).

Next, we examined the expression and localization of DMC1:interGFP in male meiocytes of wildtype and *asy1* mutants using the previously established *ASY1:RFP* reporter for staging (Yang et al. 2020b). In wildtype, DMC1:interGFP is highly expressed in both cytosol and nucleus at early prophase I (i.e., leptotene), as indicated by the initial migration of the nucleolus to one side of the nucleus and the formation of ASY1 linear threads (Yuan et al. 2025). This signal appears as numerous foci or linear stretches associated with chromosomes, alongside the diffuse signals present in the nucleoplasm (Figure 1C). This localization pattern persists till mid prophase I (i.e., mid zygotene). At late zygotene when the synapsis is largely achieved, as indicated by the removal of ASY1, the chromosome-associated DMC1:interGFP signals are decreased, with proteins primarily present in the nucleoplasm, in addition to the presence in the cytosol (Figure 1C). As meiosis further progresses to pachytene, as indicated by the complete removal of ASY1, DMC1:interGFP completely disappeared from both the cytosol and nucleus, indicating a dramatic removal of DMC1 at this stage (Figure 1C). While DMC1:interGFP localizes in both the cytoplasm and nucleus in *asy1* mutants at leptotene, we found that its chromosome association is strongly decreased compared to that in the wildtype, suggesting that ASY1 facilitates the chromosome localization of DMC1 (Figure 1D). At zygotene-like stage, this compromised chromosome association of DMC1:interGFP becomes more pronounced, exhibiting mainly a diffuse pattern within the nucleus (Figure 1D). At pachytene-like stage, evidenced by the rounded cell shape, DMC1:interGFP persists in the entire meiocytes of *asy1* mutants, in contrast to its disappearance in the wildtype. The signal remains diffuse in the nucleus with several foci present (Figure 1D). This prolonged retention of DMC1 in *asy1* persists to diplotene, where ASY3 axis is disassembled, suggesting that proper synapsis is required for the timely removal of DMC1 (Figure 1D).

### SDS is expressed at early prophase I and localizes primarily on chromatin loops

To investigate how ASY1 regulates the chromosome association of DMC1, we tested whether ASY1 and DMC1 directly interact. Both yeast two-hybrid and bimolecular fluorescence complementation (BiFC) assays showed no direct binding (Supplemental figure 2). This suggests that ASY1 likely influences DMC1 localization indirectly through modulating other factors required for its chromosome localization.

In Arabidopsis, mutations in genes involved in early recombination processes, such as DSB formation and processing, affects the chromosome recruitment of DMC1. However, the absence of ASY1 in Arabidopsis has no, or only little impact on these early recombination events (Sanchez-Moran et al. 2007; De Muyt et al. 2009), making it unlikely that ASY1 regulates DMC1 through these pathways. On the other hand, plants lacking SDS exhibit phenotypic similarities to *asy1* mutants, including defective pairing and synapsis, univalent formation, and impaired chromosome association of DMC1. Therefore, we wonder whether ASY1 regulates DMC1 localization via SDS.

To test this hypothesis, we sought to investigate whether the absence of ASY1 affects the expression and localization pattern of SDS. Since the detailed localization of SDS in plants remains poorly understood, we first characterized its expression and localization patterns during meiosis in wild-type Arabidopsis. To this end, we constructed a functional *SDS:GFP* reporter by inserting the *GFP* sequence right before the stop codon of the *SDS* genomic sequence and introgressed it along with the previously generated chromosome axis reporter ASY1:RFP into wild-type plants (Supplemental figure 3).

We observed that SDS:GFP signals start to accumulate in the nucleus of male meiocytes at late G2/early leptotene, with a diffuse localization pattern, while ASY1:RFP appears as punctate chromosomes-associated signals (Figure 2A). At leptotene, as ASY1:RFP forms the linear thread-like structures, SDS:GFP is highly expressed in the nucleus. Notably, some SDS:GFP signals clearly accumulate on the ASY1:RFP-labeled chromosomes, in addition to their presence in the nucleoplasm (Figure 2A). This localization pattern persists to zygotene, when ASY1 is progressively removed from synapsed regions (Figure 2A). At pachytene, as indicated by the complete removal of ASY1:RFP, SDS:GFP signal disappears completely (Figure 2A), suggesting SDS undergoes rapid degradation following complete synapsis, similar to the DMC1 dynamics.

**Figure 2.**
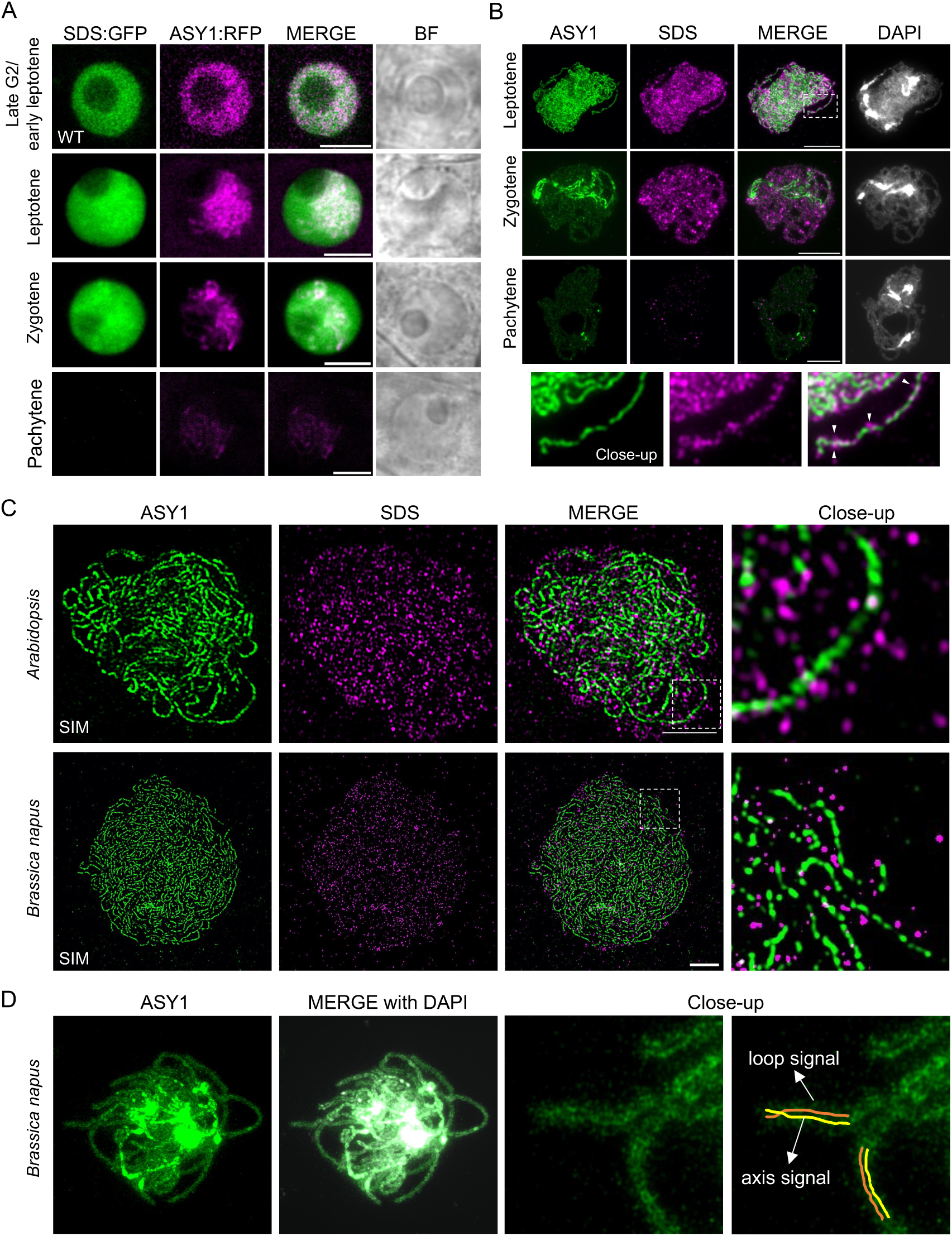
Localization pattern of SDS in wild-type Arabidopsis and *Brassica napus*. A: Live-cell confocal images displaying the expression and localization patterns of SDS:GFP during different stages of prophase I in Arabidopsis wild-type male meiocytes, with the chromosome axis highlighted by ASY1:RFP. Scale bars=5 µm. B: Co-immunostaining of ASY1 and SDS on spread chromosomes from male meiocytes of the wild-type Arabidopsis during different stages of prophase I. Images were acquired by wild-filed fluorescence microscopy. The dashed rectangle highlights the area of the close-up. Arrowheads indicate the regions where SDS localizes outside of the ASY1-marked chromosome axis. Scale bars=10 µm. C: Super-resolution imaging of the co-immunolocalization of ASY1 and SDS on spread chromosomes of the wild-type Arabidopsis and *Brassica napus* using the structured illumination microscopy (SIM). The dashed rectangles highlight the areas of the close-up. Scale bars=5 µm. D: Immunostaining of the residual ASY1 after synapsis-induced removal on spread chromosomes from male meiocytes of wild-type *Brassica napus* at pachytene. ASY1 signals from the axis and DNA loops are highlighted in the close-up panel. Scale bars=5 µm.

To further analyze the chromosome localization of SDS, we performed a co-immunostaining for SDS and ASY1 on chromosome spreads from male meiocytes. In wildtype, SDS is associated with chromosomes from early leptotene to zygotene and forms numerous foci in comparison to the linear signals of ASY1 axis (Figure 2B). In pachytene when ASY1 is fully removed from synapsed chromosomes, SDS signals become largely undetectable, consistent with the observation in live male meiocytes using the *SDS:GFP* reporter (Figure 2B). Note that no SDS signal was detected in *sds* mutants, confirming the specificity of the SDS antibody (Supplemental figure 4).

Interestingly, the signal of SDS foci seems broader than the ASY1-labeled axis from these images taken by a wide-field fluorescence microscope, indicating that some SDS proteins may localize outside of the chromosome axis, possibly on the chromatin loops (Figure 2B, close-up panel). To examine this further, we used structured illumination microscopy (SIM) providing super-resolution images. While some SDS signals co-localized with the ASY1-labeled axis, the majority formed foci surrounding the axis (Figure 2C).

To further validate these findings, we analyzed the localization of SDS (BnaSDS) in *Brassica napus*, an allotetraploid oilseed crop with larger meiocytes than in Arabidopsis, providing an advantage for studying chromosome localization of meiotic regulators. The immunolocalization of BnaSDS, using both the wide-field fluorescence microscope and super-resolution SIM, revealed a similar expression and localization pattern as that in Arabidopsis, with many BnaSDS foci surrounding the ASY1-marked axis (Figure 2C, Supplemental figure 5A).

Collectively, these results suggest that SDS, in addition to its presence in the nucleoplasm, is a chromosome-associated protein that primarily localizes on the chromatin loops organized by the chromosome axis.

### Chromosome association of DMC1 *in vivo* is reduced in the absence of SDS

Previous immunostaining studies reveal that DMC1 is nearly undetectable on chromosomes in *sds* mutants (De Muyt et al. 2009). We confirmed this finding through immunostaining of male meiocytes from *sds* mutants (Figure 3A, B), suggesting that SDS affects either the chromosome loading or protein stability of DMC1, or both. To address this question, we examined the expression and localization of DMC1 in live meiocytes of *sds-2* mutants harboring the *DMC1:interGFP* reporter.

**Figure 3.**
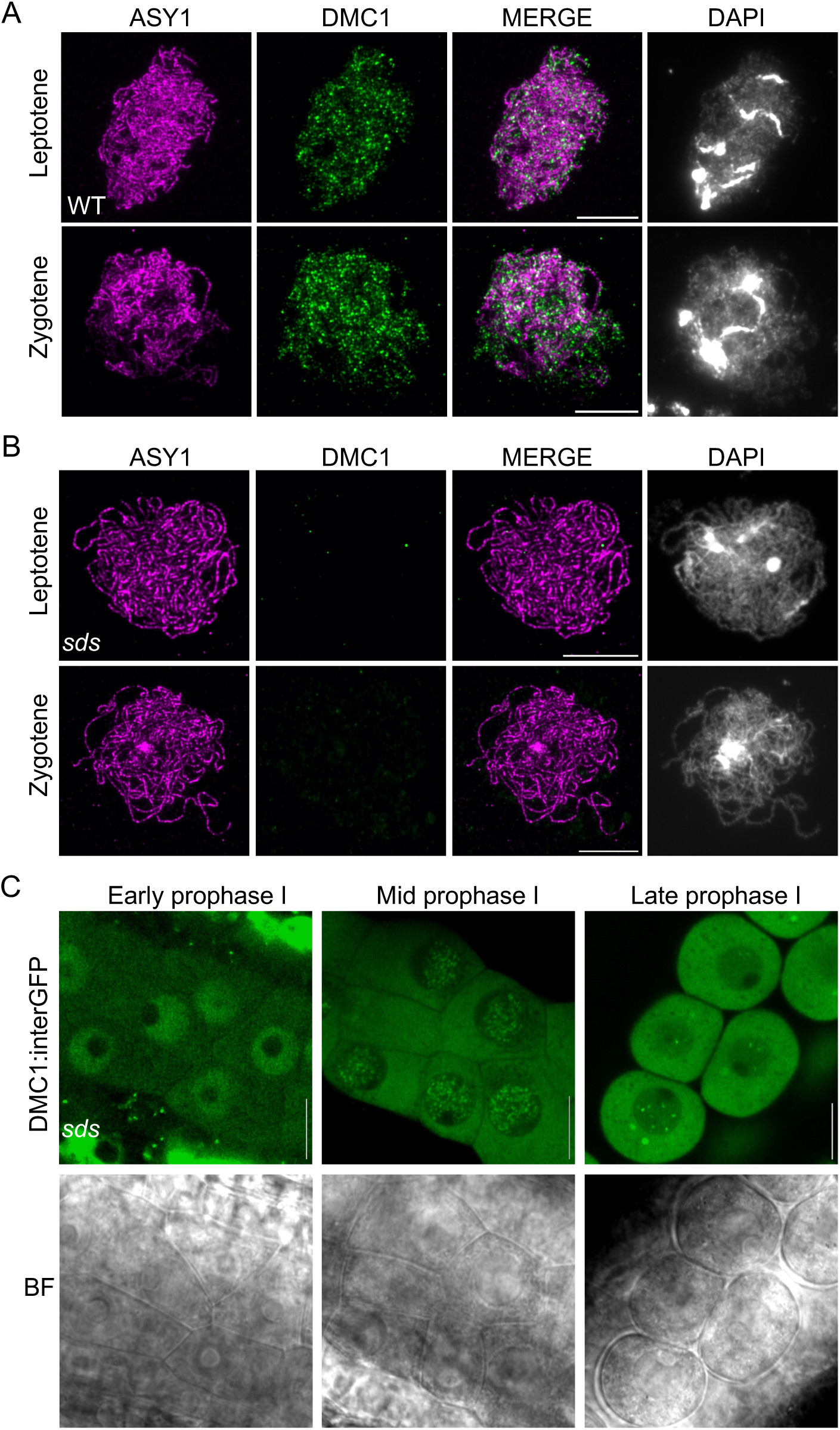
Chromosome association of DMC1 reduced is in the absence of SDS. A: Co-immunostaining of ASY1 and DMC1 on spread chromosomes from male meiocytes of the Arabidopsis wildtype (A) and *sds* mutants (B). Scale bars=10 µm. C: Live-cell confocal images showing the expression and localization patterns of DMC1:interGFP in male meiocytes of *sds* mutants. Scale bars=10 µm.

While DMC1:interGFP was observed to be present in both the cytoplasm and nucleus in *sds* mutants similar to the wildtype, we could not observe any chromosome-associated foci at early prophase I (e.g., leptotene). Instead, only diffuse GFP signals present in the nucleus, suggesting that SDS is required for the efficient loading of DMC1 to chromosomes (compare Figure 3C to Figure 1C). As meiosis further progresses to mid prophase I (zygotene-like stage), although some foci and short stretch-like signals appeared to be chromosome-associated, the overall signal within the nucleus was obviously weaker, with a substantial proportion of DMC1:interGFP remaining in the nucleoplasm (compare Figure 3C to Figure 1C). This contrasts with the strong accumulation of DMC1:interGFP on chromosomes in wild-type meiocytes (Figure 1B).

Since no chromosome-associated DMC1 could be detected in immunostaining of *sds* mutants (Figure 3B) (De Muyt et al. 2009), our results suggest that in the absence of SDS, chromosome loading/association of DMC1 becomes reduced, yet not completely abolished. In addition, at late prophase I, evidenced by the rounded shape of meiocytes, while DMC1 disappears in the wildtype (Figure 1B), the DMC1:interGFP signals remained persistent in the entire meiocytes of *sds* mutants, exhibiting only a diffuse pattern in the nucleus similar to that in *asy1* mutants (Figure 3C).

### Chromosome recruitment of SDS is strictly dependent on ASY1

Given the importance of SDS for DMC1 loading, we wonder whether ASY1 regulates DMC1 via SDS. To test this hypothesis, we studied the expression and localization of SDS in *asy1* mutants using both the live-cell imaging of the *SDS:GFP* reporter and immunostaining. The previously generated reporters *ASY3:RFP* or *ASY1:RFP* were used to mark chromosomes (Yang et al. 2020b).

Strikingly, in male meiocytes of *asy1* mutants, we found that SDS:GFP signals are present only in a diffuse pattern, in contrast to its high accumulation on chromosomes in wildtype at mid prophase I (Figure 4A). Consistently, no SDS signal could be observed on chromosomes in the immunostaining analysis of *asy1* mutants (Figure 4B). These observations suggest that the recruitment of SDS to chromosomes requires ASY1.

**Figure 4.**
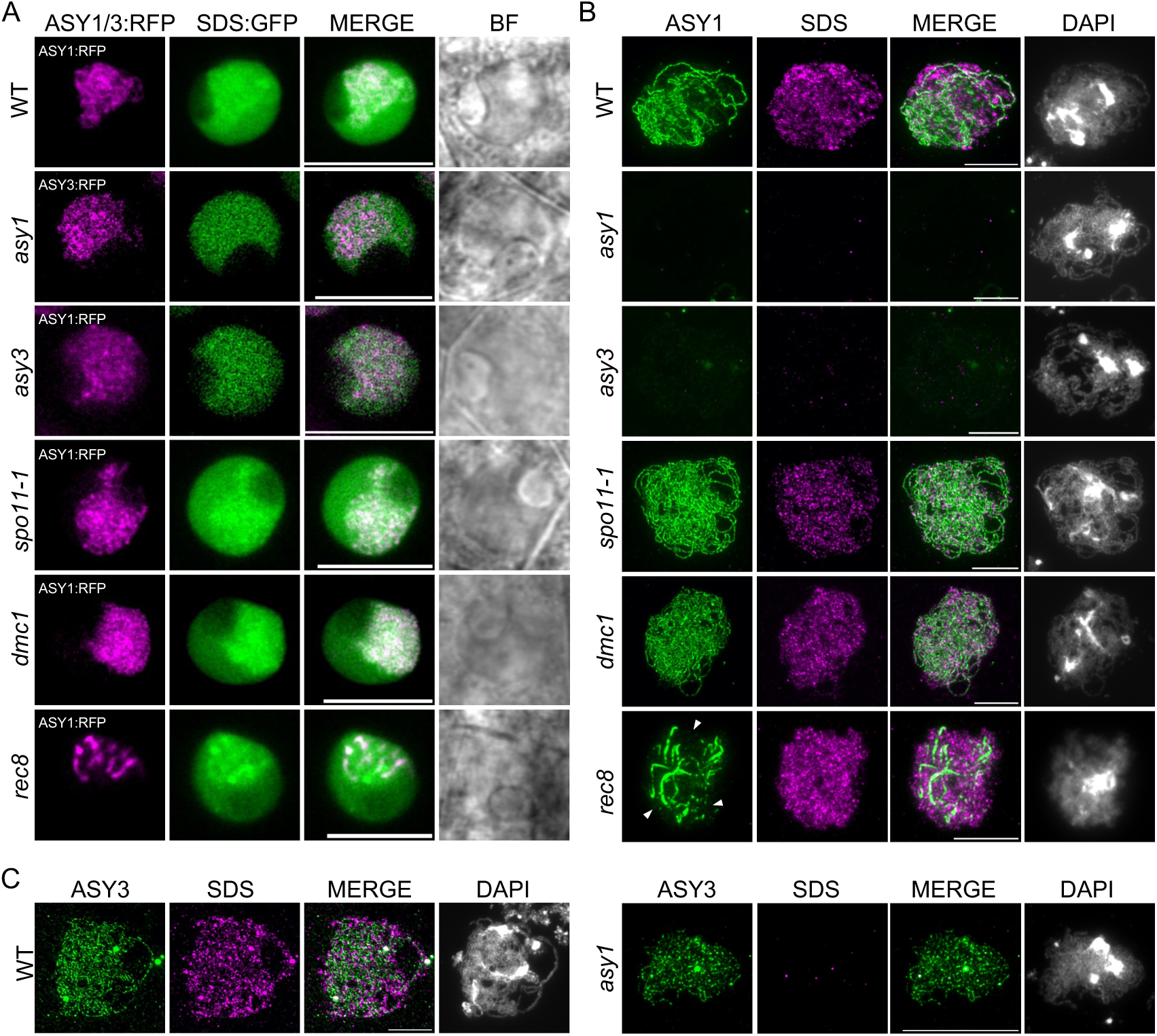
Chromosome localization of SDS in various Arabidopsis meiotic mutants. A: Live-cell confocal images showing the localization patterns of SDS:GFP in male meiocytes of *asy1*, *asy3*, *spo11-1*, *dmc1*, and *rec8* mutants, with chromosome axes highlighted by ASY1:RFP or ASY3:RFP. Scale bar = 5 µm. B: Co-immunostaining of ASY1 and SDS on spread chromosomes from male meiocytes of *asy1*, *asy3*, *rec8*, *spo11-1*, *dmc1*, and *rec8* mutants. Arrowheads indicate the fraction of ASY1 that localizes on chromosomes independent of REC8, albeit not forming bright linear axial signals. Scale bars=10 µm. C: Co-immunostaining of ASY3 and SDS on spread chromosomes from male meiocytes of the wildtype and *asy1* mutants. Scale bars=10 µm.

To determine the specificity of this dependency of SDS on ASY1, we examined the SDS localization in mutants of various genes involved in DSB formation and repair, sister chromatid cohesion, and synapsis, using both the *SDS:GFP* reporter and immunostaining. Unlike the diffuse nuclear pattern of SDS:GFP in *asy1* mutants, we found that the intensities of the chromosome-associated SDS:GFP signals in *spo11-1* and *dmc1* mutants are comparable to that in wildtype, which is further confirmed by immunostaining (Figure 4A, B). In *asy3* mutants, where the chromosome localization of ASY1 is strongly compromised, as expected, the chromosome association of SDS is largely lost (Figure 4A, B). Given that the formation of ASY3 axis is not affected in *asy1* mutants (Figure 4C) (Ferdous et al. 2012), this impaired localization of SDS in *asy3* further supports the necessity of ASY1 for SDS recruitment.

Unexpectedly, although the continuous linear structures of ASY1 axis is disrupted by the loss of REC8, the chromosome localization of SDS is not obviously affected in *rec8* mutants, as revealed by both the live-cell imaging and immunostaining (Figure 4A, B). In *rec8*, the assembly of ASY1 into linear axis structure is compromised, but a fraction of ASY1 could localizes on most chromosomes independent of REC8 (Figure 4B, arrowheads). This likely suggests that even the partial, REC8-independent loading of ASY1 on chromosomes is sufficient for SDS recruitment. Additionally, we also performed immunostaining in other mutants defective in DSB formation and repair, i.e., *atm*, *prd1*, and *mnd1*, which reveals no obvious changes in SDS chromosome association (Supplemental figure 6).

Together, these findings support that the chromosome association of SDS is strictly dependent on ASY1, but not on other key meiotic events such as DSB formation and repair. This dependency explains the impaired chromosome association of DMC1 in *asy1* mutants and substantiates that ASY1 regulates DMC1 loading through recruiting SDS. But as mentioned above, the diffuse SDS signal within in the nucleus that may be weakly associated with chromosomes is likely sufficient to promote the residual localization of DMC1 in *asy1* mutants.

Notably, the fact that ASY1 is largely removed from the synapsed chromosomes, where, however, the accumulation of SDS persists, suggests that the residual ASY1 present on both the axis and chromatin loops is sufficient for SDS recruitment (Figure 2D), further supporting the primary localization of SDS on the loop-arrays.

### ASY1 recruits SDS onto chromosomes through direct binding

Next, we sought to understand how ASY1 regulates the chromosome association of SDS. Given the strict dependency of SDS on ASY1, we wonder whether ASY1 directly binds to SDS. To test this, we performed the split-luciferase complementation assays in *N. benthamania* leaves, which reveals a direct interaction between ASY1 and SDS (Figure 5A).

**Figure 5.**
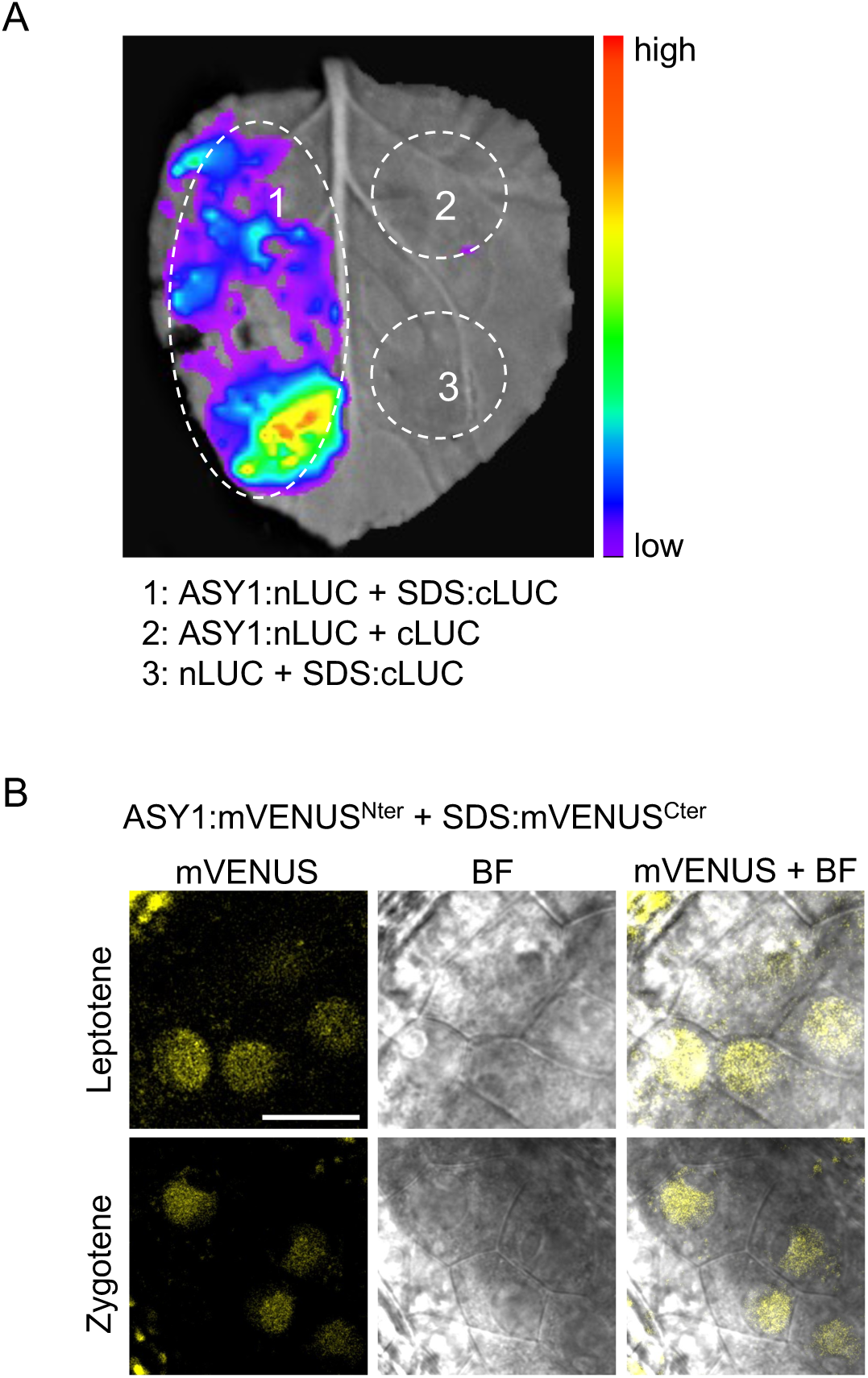
ASY1 interacts with SDS. A: Split-luciferase complementation assay for the interaction of ASY1 with SDS using *N. Benthamiana* leaves. ASY1 and SDS were fused with the N- or C-terminal parts of the luciferase, respectively. B: *In planta* BiFC assay for the interaction of ASY1 with SDS in Arabidopsis male meiocytes of wildtype. Confocal images were acquired from the anthers of the plant transformed with two complementary parts of the mVENUS protein in fusion with ASY1 (ASY1:mVenus^Nter^) and SDS (SDS:mVenus^Cter^). The mVENUS signal within the nuclei of male meiocytes indicates the interaction between ASY1 and SDS.

To further validate the interaction of ASY1 with SDS, we performed the biomolecular fluorescence complementation assays *in planta*. We generated reporter constructs for ASY1 (*PRO_ASY1_:ASY1:mVenus^Nter^*, referred to as *ASY1:mVenus^Nter^*) and SDS (*PRO_SDS_:SDS:mVenus^Cter^*, referred to as *SDS:mVenus^Cter^*), in which ASY1 and SDS were fused with the N-terminal and C-terminal fragments of the mVenus fluorescent protein, respectively. These constructs were co-transformed into Arabidopsis wild-type plants, and the presence of fluorescence signals in male meiocytes of the plants harboring both constructs was examined using confocal microscopy. As a result, in plants co-expressing ASY1:mVenus^Nter^ and SDS:mVenus^Cter^proteins, we observed complemented mVenus signals emanating mainly from the chromosomes. In contrast, the control combinations of ASY1:mVenus^Nter^ with NLS- mVeuns^Cter^ (*PRO_RPS5A_:NLS-mVenus^Cter^*) and of NLS-mVenus^Nter^ (*PRO_RPS5A_:NLS- mVenus^Nter^*) with SDS:mVenus^Cter^ did not show any obvious signals (Figure 5B, Supplemental figure 7). These results demonstrate the direct binding between ASY1 and SDS.

### Defective ASY1 removal leads to a prolonged presence of SDS and DMC1 on chromosomes

The dependency of SDS on ASY1 for its chromosome localization, together with the fact that ASY1 is largely removed from synapsed chromosomes, promoted us to investigate the effect of impaired ASY1 removal on SDS localization. To address this, we utilized *zyp1* mutants in Arabidopsis and *Brassica napus*, in which the transverse filaments of the SC are defective and ASY1 removal is abolished (Capilla-Pérez et al. 2021; France et al. 2021; Yang et al. 2022; Zhang et al. 2025).

The co-immunostaining of SDS and ASY1 revealed that, unlike in wild-type Arabidopsis and *Brassica napus* plants, where SDS disappears from synapsed chromosome at pachytene, SDS persists on chromosomes at pachytene-like stage in both *Atzyp1* and *Bnazyp1* mutants (compare Figure 2B to Figure 6A, Supplemental figure 5B). Given this extended presence of SDS, we wondered whether this would affect DMC1 dynamics. Indeed, while DMC1 is barely detected on chromosomes at pachytene in wild-type Arabidopsis plants, it remains highly abundant on chromosomes at pachytene-like stage in *Atzyp1* mutants (compare Figure 1A to Figure 6B). A similarly extended stay of DMC1 on chromosomes was also observed in *Bnazyp1* mutants (Supplemental figure 8).

**Figure 6.**
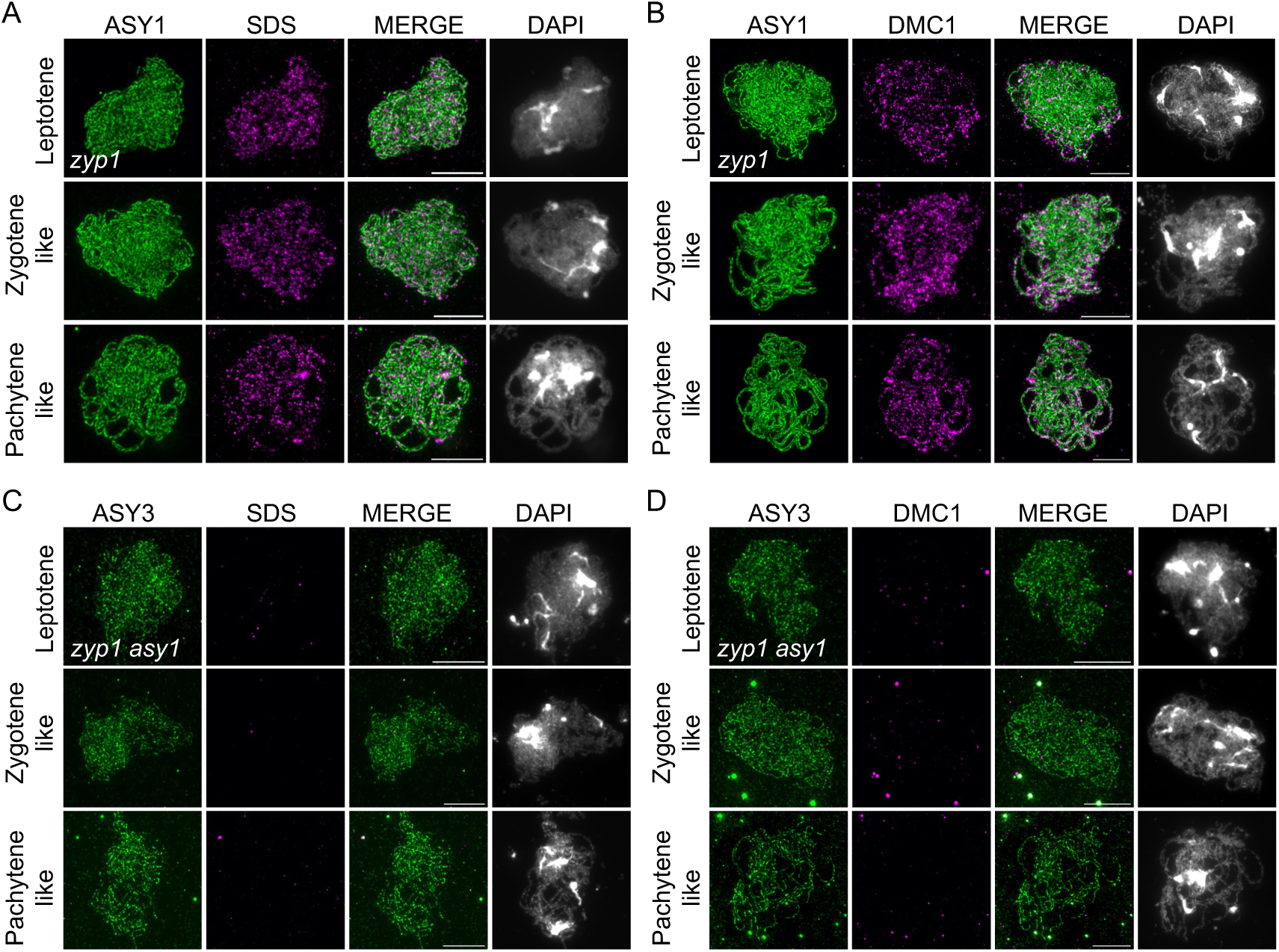
Defective ASY1 removal from the chromosome axis results in a prolonged presence of SDS and DMC1. A-B: Co-immunostaining of ASY1 and SDS (A) or DMC1 (B) on spread chromosomes from male meiocytes of the Arabidopsis *zyp1* mutants during different stages of prophase I. Scale bars=10 µm. C-D: Co-immunostaining of ASY3 and SDS (C) or DMC1 (D) on spread chromosomes from male meiocytes of the Arabidopsis *zyp1 asy1* double mutants during different stages of prophase I. Scale bars=10 µm.

To determine whether the prolonged presence of SDS and DMC1 was dependent on ASY1, we analyzed Arabidopsis *zyp1 asy1* double mutants. In these mutants, the localization of SDS and DMC1 resembles the compromised patterns observed in *asy1* mutants (Figure 6C and D). These results further substantiate the role of ASY1 in recruiting SDS, which in turn ensures proper DMC1 loading onto chromosomes.

### The closure motif of ASY1 is required for the chromosome recruitment of SDS and DMC1

Previous studies have demonstrated that the closure motif, a short peptide at the C-terminus of ASY1 that binds to its own HORMA domain, plays a crucial role in ASY1 nuclear targeting and subsequent chromosome assembly (Yang et al. 2020a; Ur and Corbett 2021). This function is mediated through an interplay between the closure motif and the AAA+ ATPase PCH2, which promotes the conformational changes of ASY1, shifting it from a self-closed state to an ‘open’ state (West et al. 2018; Yang et al. 2020a; Ur and Corbett 2021). The dependency of ASY1 localization on PCH2 is bypassed by deleting the closure motif and substituting its function in the nuclear import of ASY1 with a SV40 nuclear localization signal (NLS), as demonstrated by the wildtype-like localization of the ASY1^Δclosure^-NLS:GFP version (*PRO_ASY1_:ASY1^1-570^-NLS:GFP*, called *ASY1^Δclosure^-NLS:GFP*) in *pch2* mutants (Yang et al. 2020a). However, this ASY1 variant, lacking the closure motif, fails to complement, even not partially, the fertility and meiotic defects of *asy1* mutants (Yang et al. 2020a).

To determine whether the functional deficiency of ASY1 in the absence of the closure motif (ASY1^Δclosure^-NLS:GFP) stems from a misregulation of the recombination machinery components, we investigated whether the deletion of the closure motif affects the chromosome recruitment of SDS and DMC1. To this end, we examined the localization of SDS and DMC1 in our previously generated allele of *ASY1^Δclosure^- NLS:GFP* (*asy1*) (Yang et al. 2020a). In *asy1* mutants expressing ASY1^Δclosure^- NLS:GFP, ASY1 localization on chromosomes remained comparable to the wild-type version (Figure 7A), as previously reported (Yang et al. 2020a). However, the chromosome recruitment of SDS in *ASY1^Δclosure^-NLS:GFP* (*asy1*) was severely compromised, resembling that in *asy1* mutants (Figure 7A). Correspondingly, the chromosome localization of DMC1 was strongly impaired (Figure 7B). These results could explain the inability of ASY1^Δclosure^-NLS:GFP to rescue *asy1* mutants, and suggest that the presence of closure motif is critical for ASY1 to recruit SDS to ensure DMC1 loading, thereby promoting the interhomolog pairing and recombination.

**Figure 7.**
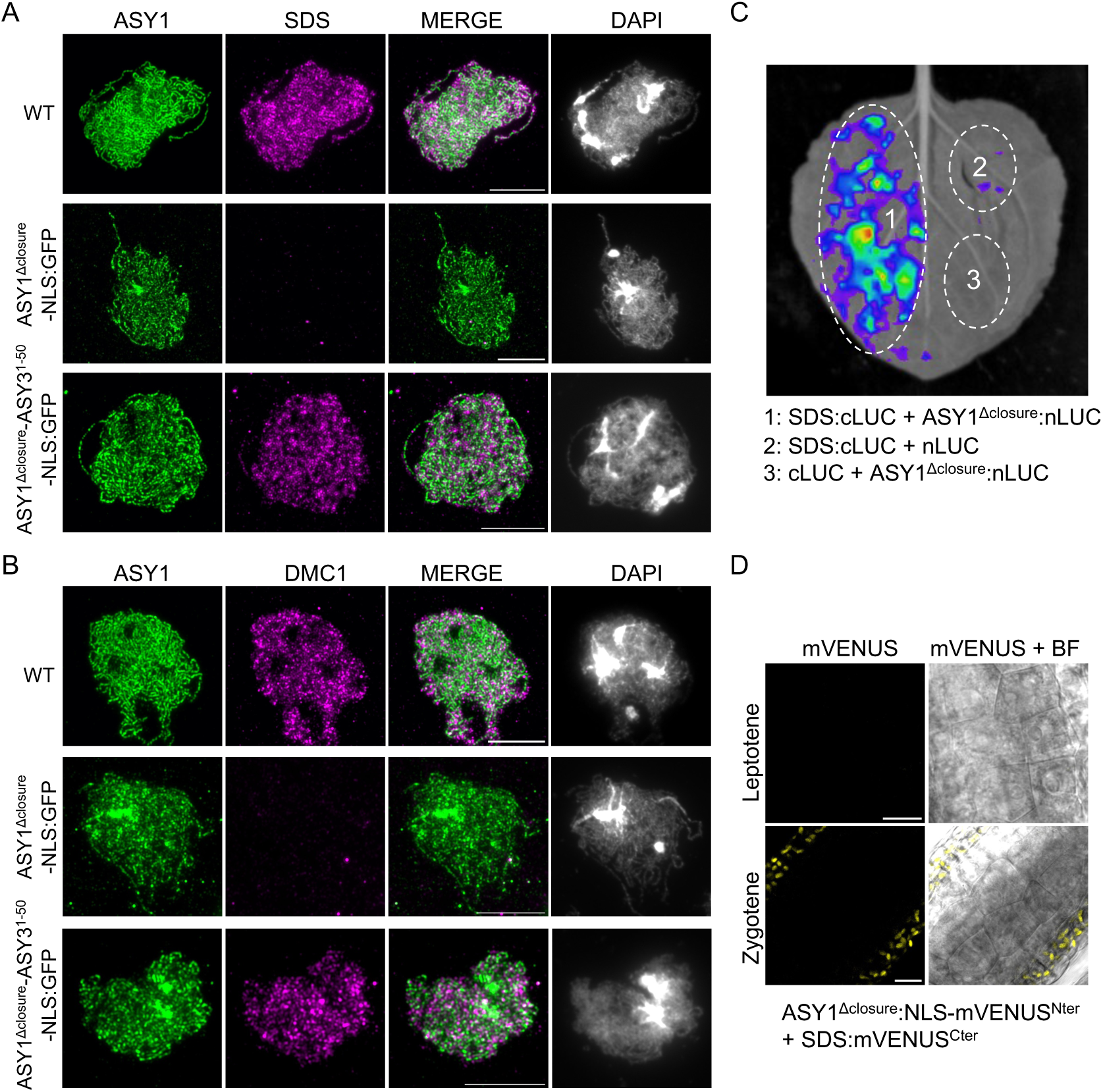
The closure motif is required for SDS and DMC1 recruitment onto chromosomes. A-B: Co-immunostaining of ASY1 and SDS (A) or DMC1 (B) on spread chromosomes of male meiocytes from wildtype, *ASY1^Δclosure^-NLS:GFP* (*asy1*), and *ASY1^Δclosure^-ASY3^1-50^-NLS:GFP* (*asy1*) mutant plants. Scale bars=10 µm. C: Split-luciferase complementation assay for the interaction of ASY1^Δclosure^ with SDS using *N. Benthamiana* leaves. ASY1^Δclosure^ and SDS were fused with the N- or C-terminal parts of the luciferase, respectively. D: *In planta* BiFC assay for the interaction of ASY1^Δclosure^ with SDS in Arabidopsis male meiocytes of wildtype. Confocal images were acquired from the anthers of the plant transformed with two complementary parts of the mVENUS protein in fusion with ASY1 (ASY1^Δclosure^:mVenus^Nter^) and SDS (SDS:mVenus^Cter^).

To explore why ASY1 lacking the closure motif fails to recruit SDS, we tested whether the absence of the closure motif affects the physical interaction between ASY1 and SDS. While a split luciferase complementation assays in *N. benthamania* leaves showed that this interaction is not affected when the closure motif is absent (Figure 7C), the interaction observed between ASY1 and SDS was lost in the *in planta* BiFC assay when an ASY1 variant without its closure motif was tested (Figure 7D). These contradictory results possibly suggest that the localization defect of SDS in *ASY1^Δclosure^-NLS:GFP* (*asy1*) is not due to the absence of the closure motif *per se* but may depend on the proper conformational state of the axis-bound ASY1.

To explore the function of the closure motif further, we utilized a previously generated substitution-of-function version of ASY1, in which ASY1’s closure motif is replaced with the HORMA domain interacting sequence of ASY3 (also referred to as a closure motif), along with the SV40 NLS to compensate for the nuclear targeting function of ASY1’s closure motif (*PRO_ASY1_:ASY1^1-570^-ASY3^1-50^-NLS:GFP*, called *ASY1^Δclosure^-ASY3^1-50^-NLS:GFP*) (Yang et al. 2020a).

We first assessed the localization of ASY1^Δclosure^-ASY3^1-50^-NLS:GFP in *asy1* mutant background and found that its chromosome association is comparable to the wild-type version of ASY1 (Figure 7A). Remarkably, we found that the chromosome localization of SDS in *ASY1^Δclosure^-ASY3^1-50^-NLS:GFP* (*asy1*) was restored compared to that in *ASY1^Δclosure^-NLS:GFP* (*asy1*) (Figure 7A). Similarly, the chromosome loading of DMC1 was also restored to the wild-type level (Figure 7B). This restoration of the chromosome recruitment of SDS and DMC1 by ASY1^Δclosure^-ASY3^1-50^-NLS:GFP was further supported by the complete rescue of the fertility of *asy1* mutants carrying this construct (Supplemental figure 9).

Taken together, these results demonstrate that the closure motif of ASY1 is required for the chromosome recruitment of SDS and DMC1, likely by ensuring a proper conformational state of ASY1 when bound to the chromosome axis (see more in the discussion).

### ASY1 acts upstream of the FIGL1-FLIP complex in regulating the chromosome localization of DMC1

The dynamic chromosome localization of DMC1 is regulated by the interplay between positive and negative regulators (Emmenecker et al. 2023). Previous studies have shown that the negative regulator FIGL1-FLIP complex represses the crucial step of stand invasion in homologous recombination, limiting crossover formation by inhibiting the chromosome association of DMC1 and RAD51 (Girard et al. 2015; Fernandes et al. 2018). Notably, the depletion of either *FIGL1* or *FLIP* can largely restore the chromosome localization defect of DMC1 in *sds* mutants, leading to a partial rescue of the crossover/bivalent formation and plant fertility (Girard et al. 2015; Fernandes et al. 2018). In our study, we identified ASY1 as a positive regulator of DMC1 by mediating the chromosome recruitment of SDS. This raises the question of whether mutating *FLIP* or *FIGL1* could also rescue the chromosome localization defect of DMC1 in *asy1* mutants.

To answer this question, we performed an epistasis analysis between *ASY1* and *FIGL1*/*FLIP* regarding their role in regulating DMC1. We carried out immunolocalization of DMC1 in double (*asy1 figl1*, *asy1 flip*) and triple (*asy1 flip sds*) mutants, in combination with staining of the chromosome axis protein ASY3 to mark chromosomes (Figure 8A). We found that while DMC1 focus formation on prophase I chromosomes was largely restored in both *sds figl1* (313.60 ± 74.60, n=15 cells vs 1.82 ± 2.98 in widetype, n=17 cells) and *sds flip* (238.58 ± 71.53, n=31 cells) double mutants, it remained compromised in both *asy1 figl1* (56.90 ± 78.23, n=28 cells) *and asy1 flip* (45.48 ± 72.41, n=40 cells), similar to that in *asy1* single mutant (33.30 ± 74.37, n=20 cells) (Figure 8A, B). This suggests that ASY1 functions upstream of FIGL1-FLIP complex in regulating DMC1 localization. In support of this conclusion, further depleting ASY1 in *sds flip* impaired the chromosome localization of DMC1, resembling the pattern seen in *asy1* mutants (Figure 8A, B).

**Figure 8.**
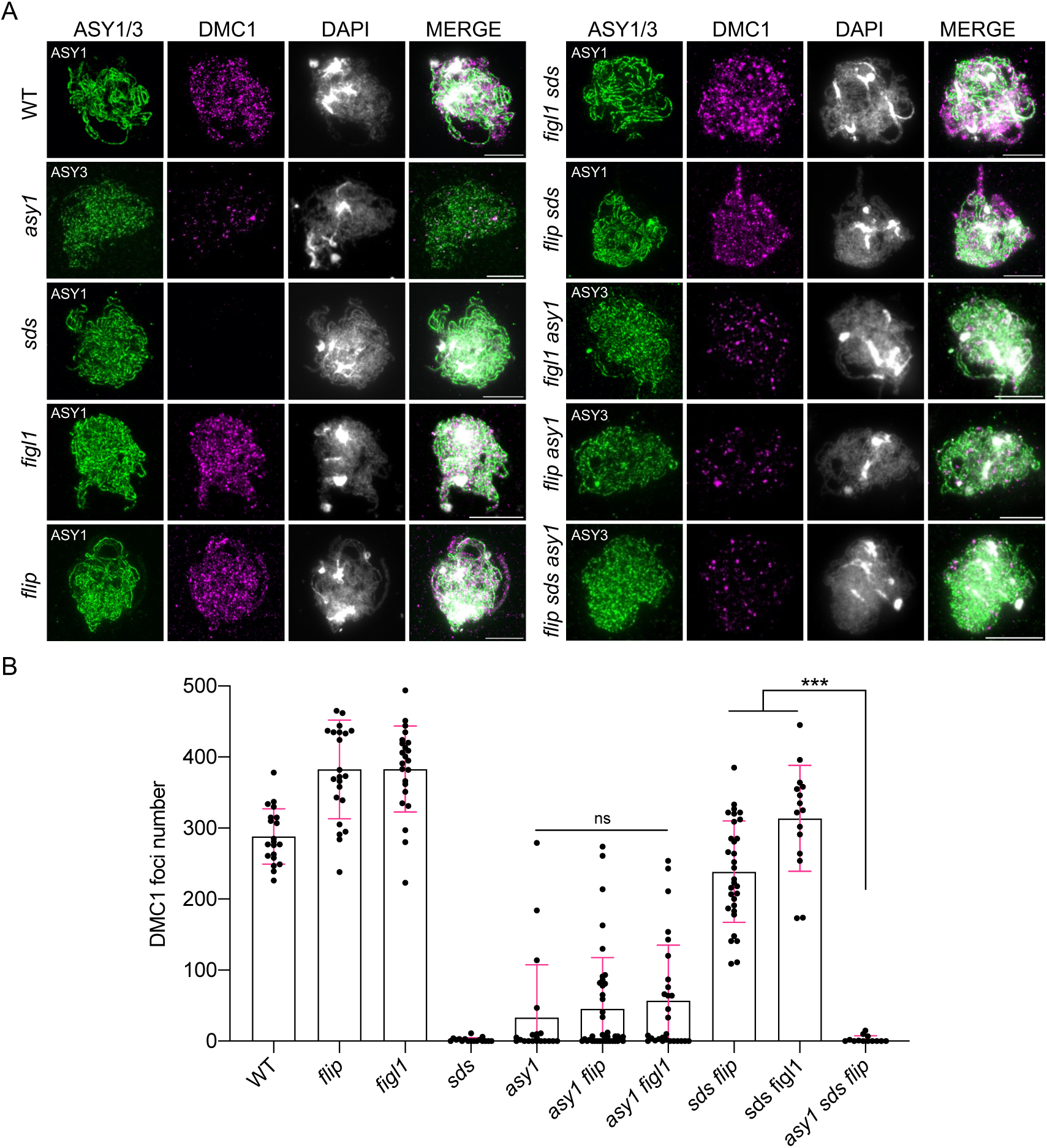
ASY1 functions upstream of the FIGL1-FLIP complex for DMC1 localization. A: Co-immunostaining of ASY1 or ASY3 with DMC1 on spread chromosomes of male meiocytes from wildtype, *asy1*, *sds*, *figl1*, *flip*, *figl1 sds*, *flip sds*, *figl1 asy1*, *flip asy1*, and *flip sds asy1* mutant plants at early prophase I (e.g., late leptotene or zygotene). Scale bars=10 µm. B: Statistical analysis of the numbers of DMC1 foci shown in (A). Asterisks indicate significant difference (Tukey’s multiple comparisons test, p<0.001) and ns indicates no significant difference.

## Discussion

The chromosome axis is a critical structural and regulatory platform for meiotic recombination coordinating different steps from DSB formation, over homology search, to crossover formation. The key role of the axis in homologous recombination is to recruit and/or anchor many factors of the recombination machinery including the recombinases themselves. However, the mechanisms by which the axis coordinates the recombinase dynamics remain poorly understood. The axial component ASY1 in Arabidopsis has been shown to play a role in recruiting and/or stabilizing the chromosome localization of DMC1, yet by an unclear mechanism (Sanchez-Moran et al. 2007). Our study reveals that ASY1 ensures accurate DMC1 loading onto chromosomes by directly recruiting the plant-specific cyclin SDS, a positive regulator of DMC1 localization. Furthermore, we demonstrate that ASY1 acts upstream of the FIGL1-FLIP negative regulatory complex, positioning it as a key coordinator of DMC1 localization dynamics. These findings advance our understanding of how chromosome axis architecture interacts with the recombination machinery to ensure homologous pairing and crossover formation.

### ASY1 bridges chromosome axis architecture and recombinase recruitment

The chromosome axis has long been shown to act as a scaffold for recombination factors (Sanchez-Moran et al. 2008; Lam and Keeney 2015; Zickler and Kleckner 2015, 2023), but the direct mechanistic link between the axis and recombinase loading have remained elusive. Our discovery that Arabidopsis ASY1 directly binds to SDS, a key promoter of DMC1 loading, provides a missing link. We show that SDS is present both in the nucleoplasm and on chromosomes where it forms numerous foci. In the absence of ASY1, the recruitment of SDS onto chromosomes is compromised, which in turn affects the chromosome loading of DMC1. Conversely, a persistent stay of ASY1 on axis, as seen in both Arabidopsis and *Brassica napus zyp1* mutants, leads to a prolonged stay of SDS and thus DMC1 on chromosomes, supporting the observation in several species that the SC assembly seem to limit the DSB formation and repair machinery to non-synapsed sections of axes where DSBs are used for homologous pairing and synapsis (Panizza et al. 2011; Thacker et al. 2014; Stanzione et al. 2016; Mu et al. 2020; Wang et al. 2023). Our results support the idea that this regulatory mechanism involve an SC-triggered depletion of meiotic HORMADs from synapsed axes (Wojtasz et al. 2009; Yang et al. 2022).

This ASY1-dependent chromosome recruitment of SDS explains why *asy1* mutants largely phenocopy *sds* mutants in their defective DMC1 loading, pairing, and bivalent formation (Azumi et al. 2002; Sanchez-Moran et al. 2007; De Muyt et al. 2009).

Nevertheless, the mutation of *SDS* results in nearly a complete abolishment of DMC1 loading, which is more severe than that in *asy1* mutants, suggesting that the rest of SDS present in nucleoplasm or those weakly associated with chromosomes in *asy1* can still promote a low level of DMC1 loading. This reduced DMC1 level could mediate the formation of crossovers observed at the chromosome ends of *asy1* mutants, in contrast to the complete abolishment of crossovers in *sds* mutants (De Muyt et al. 2009; Lambing et al. 2020a; Pochon et al. 2023). Our results support the idea that ASY1 functions as a structural rather than catalytic regulator of recombination, ensuring the spatiotemporal patterning of DMC1 activity at the axis for efficient interhomolog-biased DSB repair in Arabidopsis. A question related to this is to which extent the elevated crossover formation in the absence of SC, e.g., in *zyp1* mutants, is attributed to the prolonged stay of DMC1 and possibly other recombination proteins (Yang et al. 2022). Notably, the dependency of SDS on ASY1 for chromosome association is highly specific. While SDS localization is unaffected in mutants defective in DSB formation (*spo11-1*), sister chromatid cohesion (*rec8*), or synapsis (*zyp1*), it is completely lost in *asy1* mutants. This also supports the conclusion that ASY1’s role in SDS recruitment is not merely a passive consequence of axis assembly but involves direct molecular interaction, namely, ASY1-SDS binding is essential for DMC1 loading. This mechanism resembles the observations in yeast, mouse, and maize, where Hop1/HORMAD1/ZmASY1 (the Arabidopsis ASY1 homolog) recruits the DSB-promoting factor Mer2/IHO1/ZmPRD3 to facilitate the clustering of DSB machinery on axes, thereby enabling efficient DSB formation (Panizza et al. 2011; Stanzione et al. 2016; Kariyazono et al. 2019; Rousova et al. 2021; Wang et al. 2023; Dereli et al. 2024).

### Role of ASY1’s closure motif in SDS recruitment

The presence of closure motif confers ASY1 the ability to change conformations for its nuclear targeting and axis assembly, which is mediated by the AAA+ ATPase PCH2 (Ye et al. 2017; Yang et al. 2020a). While a previous study showed that the closure motif, when complemented with an exogenous NLS, is dispensable for ASY1’s nuclear import or axis localization (Yang et al. 2020a), we found that its deletion strongly compromises the chromosome loading of SDS and DMC1. Nevertheless, the split-luciferase complementation assays in *N. benthamania* leaves but not the in *planta* BiFC show that the interaction of ASY1 and SDS persists in the absence of the closure motif. These inconsistent observations suggest that the closure motif, through interacting with the N-terminal HORMA domain, perhaps enables a conformationally permissive state of the axis-associated ASY1, facilitated by PCH2-mediated remodelling, which is necessary for SDS recruitment. The absence of the closure motif abolishes such a state. This model is supported by the rescue of SDS and DMC1 loading when the ASY1’s closure motif is replaced with the analogous peptide motif from ASY1’s binding partner ASY3 that binds to the HORMA domain. Future study on investigating the structural basis of ASY1-SDS interaction could elucidate how the closure motif-dependent conformational changes of ASY1 enable SDS recruitment. The role of the closure motif in SDS recruitment underscores the importance of HORMAD protein dynamics in meiosis.

### ASY1 coordinate opposing regulators in DMC1 dynamics

The chromosome localization of DMC1 is tightly regulated by a tug-of-war between positive (e.g., SDS, BRCA2) and negative (e.g., FIGL1-FLIP, SMC5/6) regulators. Our epistasis analysis places ASY1 upstream of FIGL1-FLIP complex, as depleting FIGL1/FLIP fails to restore DMC1 foci in *asy1* mutants. This contrasts with *sds* mutants, where FIGL1/FLIP depletion substantially rescues DMC1 loading (Girard et al. 2015; Fernandes et al. 2018), indicating that the function of ASY1 extends beyond SDS recruitment. These observations suggest that the axis likely provides a basic context for both the positive and negative regulators of DMC1 to work on.

The fact that the chromosome recruitment of DMC1, but not SDS, is highly dependent on DSB formation, indicating that SDS likely functions to stabilizes the formation of DMC1-ssDNA filaments. The underlying mechanism of how SDS stabilizes DMC1 loading to ssDNA remains to be resolved. We propose that the axis assembly of ASY1 establishes a permissive chromatin environment for DMC1 activity by recruiting SDS to stabilize DMC1-ssDNA interaction, which counteracts the FIGL1-FLIP inhibition, e.g., SDS might exclude the FIGL1-FLIP complex from recombination sites.

In conclusion, the results we present here provide insights into the longstanding question about how the chromosome axis orchestrates meiotic recombination. We conclude that ASY1 plays dual roles in controlling DMC1 activity, namely, promoting DMC1 activity by ensuring its loading at early prophase I while restraining its excess through a synapsis-dependent removal. This positions ASY1 as a central integrator of recombination efficiency and fidelity.

## Materials and methods

### Plant Materials

The *A. thaliana* ecotype Columbia-0 (Col-0) and *Brassica napus* variety Westar were used as the wild-type reference in this study. The Arabidopsis T-DNA insertion lines SAIL_129_F09 (*sds-2*), SALK_146172 (*spo11-1-3*), SAIL_807_B08 (*rec8*), SAIL_170_F08 (*dmc1-2*), SALK_046272 (*asy1-4*), SALK_143676 (*asy3-1*), SALK_037387 (*flip-2*), SALK_058551 (*figl1-17*), SALK_006953 (*atm-2*), SALK_024703 (*prd1-2*) and SALK_110052 (*mnd1*) were obtained from the Arabidopsis sharing center in China (https://www.arashare.cn) and the European Arabidopsis Stock Centre (NASC, https://arabidopsis.info/). The *Brassica napus zyp1* mutants were created previously by CRISPR-Cas9 (Zhang et al. 2025). The Arabidopsis T-DNA insertion mutant lines together with detailed genotyping information are listed in Supplementary Table 1. Primers used for constructing plasmids and genotyping T-DNA mutants are shown in Supplementary Table 2. All plants were cultivated in growth chambers with a 16h /21 °C light and 8h/18°C dark photoperiod and 60% humidity.

### Reporter construction and plant transformation

To construct SDS reporters, the genomic fragment of SDS including the DNA sequences of 1535 bp before the start codon and of 452 bp after the stop codon of SDS were amplified using the primers of AtgSDS-attB1-F and AtgSDS-attB2-R (Supplementary Table 2). The PCR product was cloned into pDONR221 via the gateway BP reaction, creating *PRO_SDS_:SDS_pDONR221*. Next, the GFP fragments were inserted into the position right before the stop codon using a SLICE reaction, generating the *PRO_SDS_:SDS:GFP_pDONR221* entry clones. For the SLICE reaction, the primer pair of SDS-SLICE-CGFP-F with SDS-SLICE-CGFP-R was used to linearize the *PRO_SDS_:SDS_pDONR221* entry clone by PCR (Supplementary Table 2). Next, the expression cassette of *PRO_SDS_:SDS:GFP* was integrated into the destination vector *pGWB501* by the gateway LR reaction, producing *PRO_SDS_:SDS:GFP_pGWB501* reporter construct.

For the construction of DMC1:interGFP reporters, the genomic sequence of DMC1 covering the DNA sequences of 3446 bp before the start codon and of 554 bp after the stop codon was amplified using the primer pair of AtDMC1-attB1-F and AtDMC1-attB2-R (Supplementary Table 2). The PCR product was cloned into the entry vector pDONR221 via the Gateway BP reaction, producing *PRO_DMC1_:DMC1_pDONR221*. Next, the GFP sequence was inserted internally between 5 and 6 amino acids of DMC1 using the SLICE reaction, resulting in the entry vectors of *PRO_DMC1_:DMC1:interGFP_pDONR221*. For the SLICE reaction, the primer pairs of DMC1-SLICE-interGFP-F with DMC1-SLICE-interGFP-R were used to linearize *PRO_DMC1_:DMC1:interGFP_pDONR221* entry clone by PCR (Supplementary Table 2). Next, the expression cassette of *PRO_DMC1_:DMC1:interGFP* was integrated into the destination vector pGWB501 by the gateway LR reaction, producing the *PRO_DMC1_:DMC1:interGFP_pGWB501* reporter construct. All reporter constructs were transformed into *A. thaliana* wild-type and relevant mutant plants by using the *Agrobacterium tumefaciens* (GV3101)-mediated floral dipping method.

### Yeast two-hybrid assay

To assess the interaction between ASY1 and DMC1 using a yeast two-hybrid assay, the constructs of ASY1-AD and DMC1-BD were generated. The coding sequences (CDS) of ASY1 and DMC1 were amplified via PCR using primers flanked by *attB* recombination sites (ASY1-CDS-attB1-F and ASY1-CDS-attB2-R; DMC1-CDS-attB1-F and DMC1-CDS-attB2-R) (Supplementary Table 2). The resulting PCR products were cloned into the pDONR223 vector through Gateway BP recombination. Subsequently, these entry clones were integrated into the destination vectors pGADT7-GW and pGBKT7-GW via Gateway LR reaction.

For the yeast two-hybrid interaction assay, the relevant plasmid combinations were co-transformed into the AH109 *Saccharomyces cerevisiae* strain using the polyethylene glycol/lithium acetate method, following the manufacturer’s protocol (Clontech). Positive transformants were selected on synthetic dropout (SD) medium lacking leucine and tryptophan (SD/-Leu/-Trp). Interaction between tested proteins was assessed by spotting the yeast cells onto plates of triple (-Leu/-Trp/-His) and quadruple (-Leu/-Trp/-His/-Ade) synthetic dropout media. After incubating at 28 °C for three days, yeast growth was imaged to evaluate protein-protein interactions.

### Split luciferase complementation assay

For the split-luciferase complementation assays, the constructs of ASY1-nLUC, SDS-cLUC, DMC1-nLUC, ASY1-cLUC, and ASY1^Δclosure^-nLUC were generated using a dual-enzyme digestion cloning approach with the pCambia1300-nLUC and pCambia1300-cLUC vectors. All genes were driven by the 35S promoter. The resulting constructs were transformed into *Agrobacterium* strain GV3101. The agrobacteria were grown at 28 °C and harvested at OD_600_ of approximate 1.0. The bacteria pellets were resuspended in the infiltration buffer containing 10 mM MES, 100 μM acetosyringone, and 10 mM MgCl2. Equal volumes of agrobacteria for the relevant combinations were mixed, and an additional 10% volume of agrobacteria containing the P19 expression vector was added into the mixture to inhibit gene silencing. The relevant combinations of agrobacteria were injected into *N. benthamiana* leaves. The plants were kept in the dark for 12h and then put back to the normal growth condition for 36-48 hours. To evaluate the interaction, all leaves were injected with the solution of 0.3 mg/mL D-luciferin, and then, the luciferase signals were captured using the NightSHADE L985 imaging system (Berthold Technology).

### Bimolecular fluorescence complementation assay

For the *in planta* BiFC assays, we constructed the *PRO_ASY1_:ASY1:mVENUS^Nter^_pDONR221*, *PRO_SDS_:SDS:mVENUS^Cter^_pDONR221* and *PRO_ASY1_:ASY1^Δclosure^-NLS:mVENUS^Nter^_pDONR221* entry clones by replacing the GFP tag in the previously generated reporter constructs of *PRO_ASY1_:ASY1:GFP_pDONR221* (Yang et al. 2020b) and *PRO_ASY1_:ASY1^Δclosure^-NLS:GFP_pDONR221* (Yang et al. 2020a), and the here-generated *PRO_SDS_:SDS:GFP_pDONR221* with the *mVENUS^Nter^* or *mVENUS^Cter^* fragments using the SmaI restriction sites flanking the GFP tag. Subsequently, these entry clones were integrated into the destination vectors pGWB501 or pGWB601 using Gateway LR reactions, producing the *PRO_ASY1_:ASY1:mVENUS^Nter^_pGWB501*, *PRO_SDS_:SDS:mVENUS^Cter^_pGWB601*, and *PRO_ASY1_:ASY1^Δclosure^-NLS:mVENUS^Nter^_pGWB501* vectors. For the control constructs of *PRO_RPS5A_:NLS-mVENUS^Nter^_R4pGWB501* and *PRO_RPS5A_:NLS-mVENUS^Cter^_R4pGWB601*, the NLS sequence was introduced in front of the mVENUS^Nter^ and mVENUS^Cter^ fragments by PCR using primers flanked by attB1/B2 recombinaton sites. The resulting PCR products were cloned into the entry vector pDONR221 using the gateway BP reaction, resulting in the *NLS-mVENUS^Nter^_pDONR221* and *NLS-mVENUS^Cter^_pDONR221* entry vectors. Subsequently, the previously generated RPS5A promoter entry clone (*PRO_RPS5A__P4P1r-pDONR*) was combined with the *NLS-mVENUS^Nter^_pDONR221* and *NLS-mVENUS^Cter^_pDONR221* entry vectors through multisite Gateway LR reactions with the destination vectors R4pGWB501 or R4pGWB601, resulting in the *PRO_RPS5A_:NLS-mVENUS^Nter^_R4pGWB501*and *PRO_RPS5A_:NLS-mVENUS^Cter^_R4pGWB601* constructs. We then co-transformed the relevant combinations of these constructs into Arabidopsis wild-type plants via the *Agrobacterium*-mediated floral dipping method. Transgenic plants harboring both constructs were selected on ½MS medium supplemented with both hygromycin (25 mg/L) and basta (10 mg/L). Meiotic stage anthers were dissected and visualized directly under a confocal microscope with an excitation wavelength of 560-580 nm.

### Antibody generation

The polyclonal antibodies against Arabidopsis and Brassica SDS were produced by DIA-AN, Wuhan, China (https://www.dia-an.com). Briefly, the coding regions of AtSDS and BnaC08.SDS (121-345 aa) were amplified and inserted into the pET-32a vector. The corresponding recombinant proteins were produced and purified from *Escherichia coli* bacteria, which were used as antigens to immunize rats. After 3-time immunization, antibodies were purified from the antisera using antigen-based affinity purification.

### Chromosome spreads

Young flower buds were harvested and immediately fixed in the ice-cold Carnoy’s fixative solution (3:1 ethanol: glacial acetic acid, v/v) for 48 hours and then stored at -20°C for longer storage. Meiotic-stage buds, approximately 0.3-0.5mm in size, were dissected and soaked in distilled water for 10 minutes to remove the fixative solution, followed by the enzyme digestion (3% cellulose, 3% macerozyme, and 5% snailase in 50 mM citrate buffer, pH 4.5) for 1 hour at 37°C. After rinsing with water, the flower buds were transferred onto an adhesive microscope slide and squashed completely in 10-15 µl of 45% acetic acid. Next, another 10-15 µl of 45% acetic acid was added to each sample, followed by another round of smashing. Subsequently, an additional 10-15 μl of 45% acetic acid was added onto the slide, and chromosomes were spread on a 45°C hotplate by stirring the solution until it was largely evaporated. The slide was immediately washed with ice-cold fixative solution. Once the slide was air-dried, the chromosomes were stained using the VECTASHIELD antifade mounting medium containing DAPI and imaged under a SOPTOP RX50 fluorescent microscope (Sunny Optical Technology, China) equipped with a 100× oil immersion objective (Olympus, numerical aperture (NA) = 1.3) and a monochrome camera.

### Confocal microscopy for observing reporters in live male meiocytes

The *in vivo* observation of fluorescent reporter proteins in live male meiocytes was performed according to the previous description (Prusicki et al. 2019). Briefly, the fresh anthers were isolated from flower buds of approximately 0.3 to 0.5 mm and placed on a slide with about 70 µl of water. After covering the slide with a coverslip, the anthers were immediately imaged using a Leica TCS SP8 inverted confocal microscope equipped with an HC APO L 63x/0.90 W UV-I objective. The stages of meiosis were determined based on the combined criteria including cell shape, nucleolus position, and chromosome configuration as described in (Prusicki et al. 2019).

### Immunolocalization

For immunostaining on acetic acid-spreads, slides were prepared using the chromosome spread method mentioned above and then examined under a phase-contrast microscope to identify those containing the desired meiotic cells. Next, the slides were immersed in a washing jar containing 10 mM sodium citrate buffer (pH 6.8) and microwaved until the liquid was nearly boiling. They were immediately transferred to the PBST buffer (1x PBS + 1% Triton X-100) and incubated at room temperature for 10 minutes. The slides were blocked with commercial goat pre-immune serum for 2 hours at room temperature. Subsequently, primary antibodies (listed in Supplementary Table 3) diluted in goat pre-immune serum at a ratio of 1:250 were applied to the slides and incubated at 4°C for 48 hours. After three washes with PBST buffer for 5 minutes each, fluorescently labeled secondary antibodies (as detailed in Supplementary Table 3) were applied and incubated at 4°C for 24 hours. The slides were then washed three times with PBST, and the chromosomes were stained with an antifade DAPI solution (VECTASHIELD) overnight. The slides were examined under a SOPTOP RX50 fluorescent microscope (Sunny Optical Technology, China) with an 100x oil immersion objective (Olympus).

## Acknowledgements

This work was supported by the National Key Research and Development Program of China (2023YFF1000700 to CY), National Natural Science Foundation of China (32370360, 32170354 to CY), Hubei Key Research and Development Program (2023BBB173 to CY), and Fundamental Research Funds for the Central Universities (2662023ZKPY003 and 2662023PY004 to CY).

## Author contributions

C.Y. conceived this research. B.C., X.Y., Y.L., F.C., Y.Z., J.Z., M.G., and L.C. performed the experiments. B.C., Y.L., F.C., and C.Y analyzed the data. C.Y., A.S., and B.C. wrote the manuscript.

