## Supplementary figures 1-9 for "The HORMA-domain protein ASY1 recruits the plant-specific cyclin SDS to promote DMC1-mediated meiotic recombination"

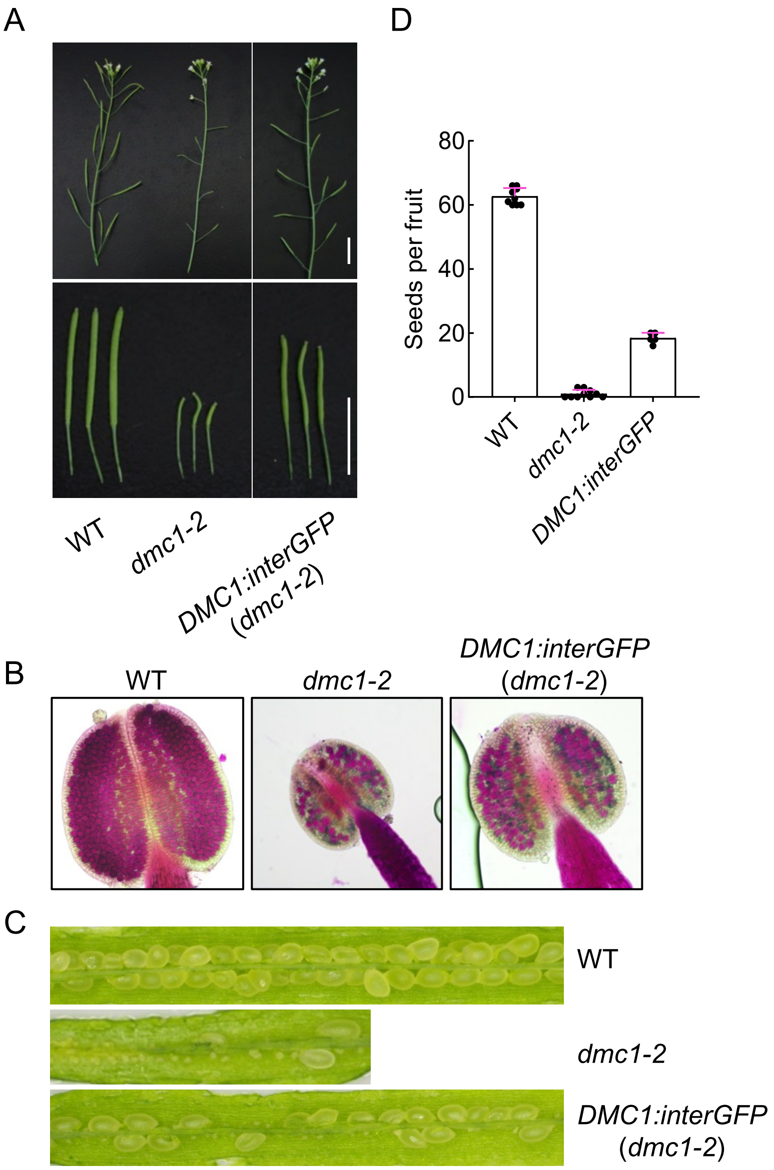


**Supplementary Figure 1. The *DMC1:GFP* reporter is partially functional.** (A) The main branches (upper panel) and siliques (low panel) of WT, *dmc1-2* and *DMC1:interGFP* (*dmc1-2*) plants. Bars: 1 cm. (B) Pollen staining of WT, *dmc1-2* and *DMC1:interGFP* (*dmc1-2*) plants. (C) The seed sets of WT, *dmc1-2* and *DMC1:interGFP* (*dmc1-2*) plants. (D) Statistical analysis of seed sets according to (C).

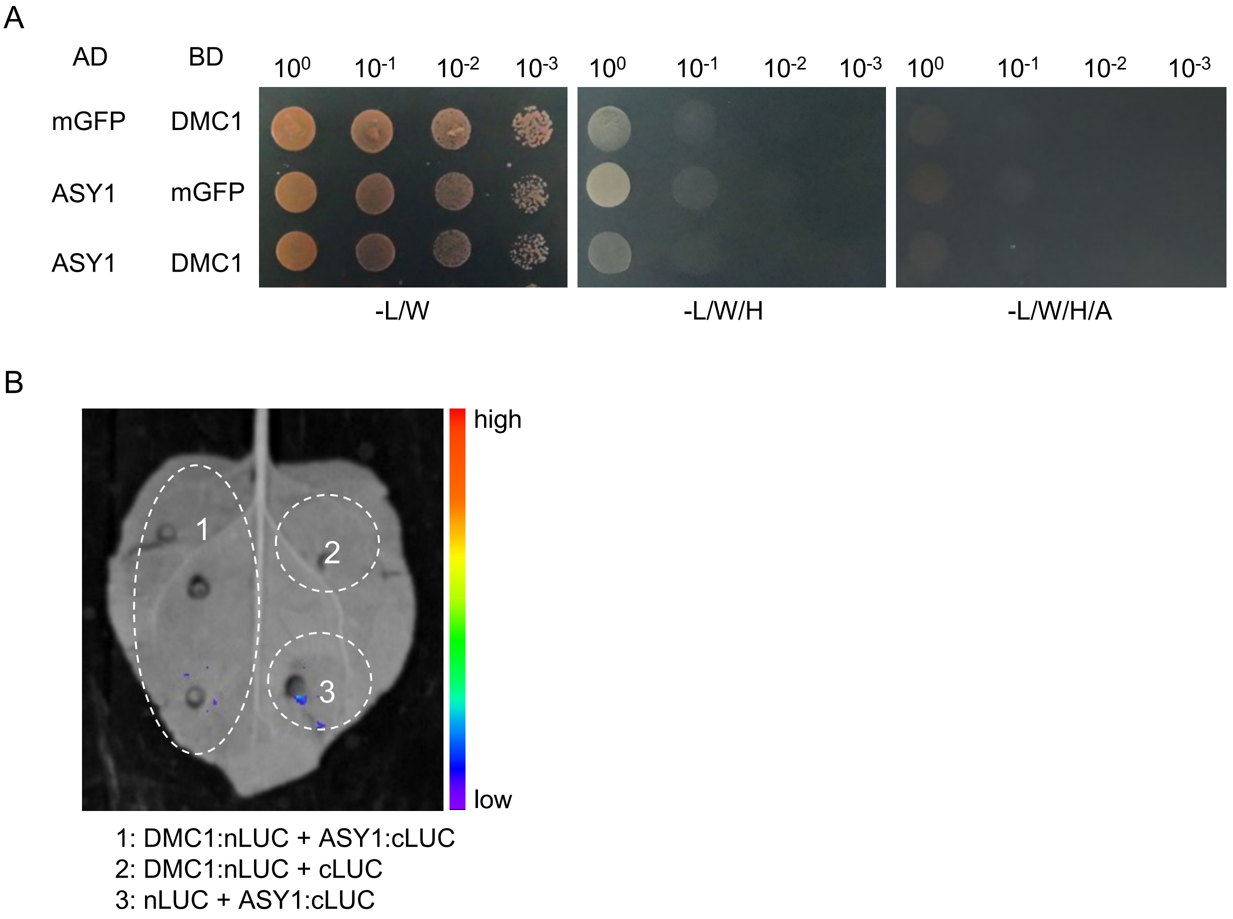


**Supplementary Figure 2. ASY1 has no direct physical interaction with DMC1**. (A) Yeast two-hybrid assay for testing the interaction between ASY1 and DMC1. Monomeric GFP (mGFP) fused with AD (activating domain) and BD (binding domain) were used as controls. (B) Split-luciferase complementation assay for the interaction of ASY1 with DMC1 using *N. Benthamiana* leaves. ASY1 and DMC1 were fused with the C- or N-terminal parts of the luciferase, respectively. The unfused forms of nLUC or cLUC were used as negative controls.

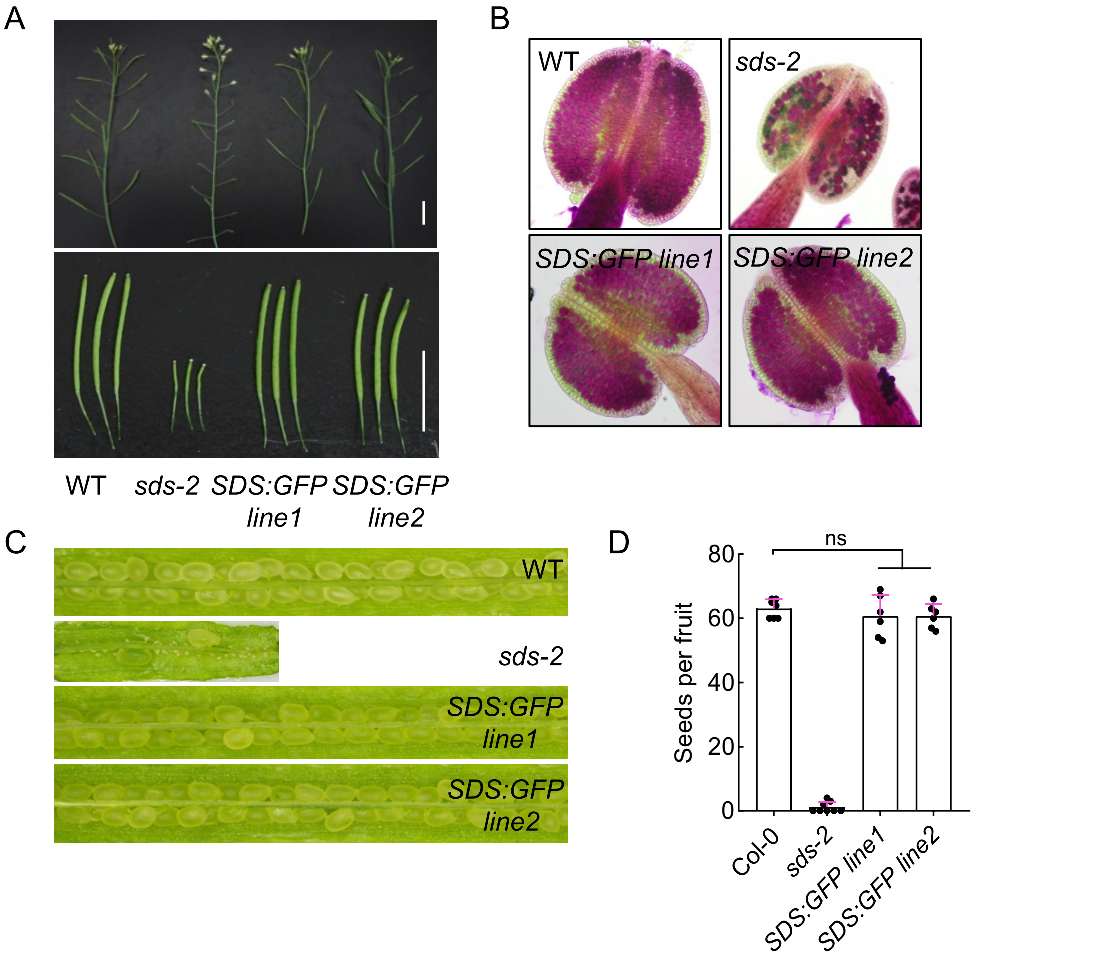


**Supplementary Figure 3. The SDS:GFP reporter is functional.** (A) The main branches (upper panel) and siliques (low panel) of WT, *sds-2*, and two lines of *SDS:GFP* (*sds-2*) plants. Bars: 1 cm. (B) Pollen staining of WT, *sds-2*, and two lines of *SDS:GFP* (*sds-2*) plants. (C) The seed sets of WT, *sds-2*, and two lines of *SDS:GFP* (*sds-2*) plants. (D) Statistical analysis of seed sets according to (C).

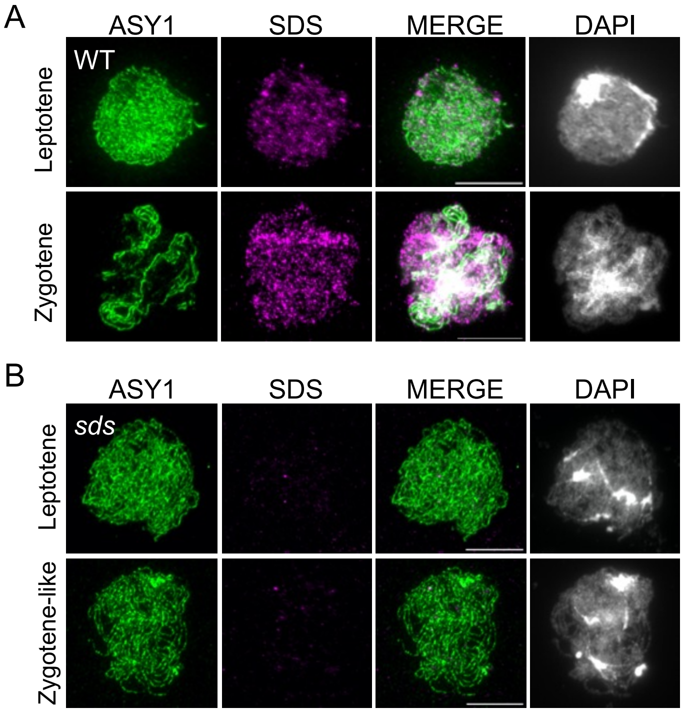


**Supplementary Figure 4. The specificity validation of SDS antibody.** (A) Co-immunostaining of ASY1 and SDS in male meiocytes of the wildtype Arabidopsis during prophase I. (B) Co-immunostaining of ASY1 and SDS in male meiocytes of *sds* mutants during prophase I. Scale bars=10 μm.

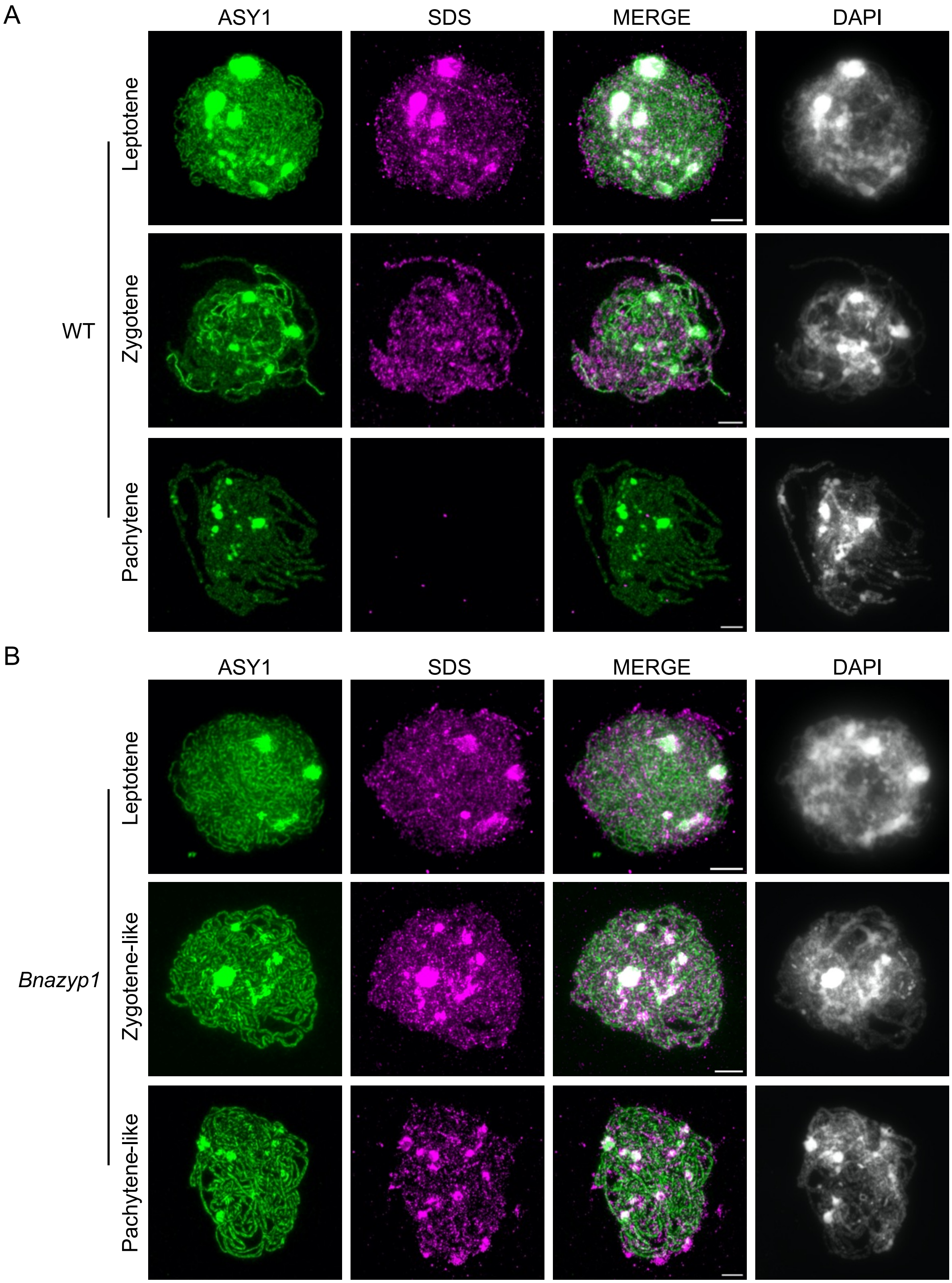


**Supplementary Figure 5. SDS exhibits prolonged stay on meiotic chromosomes in *Brassica napus zyp1* mutants.** (A) Co-immunolocalization of SDS and ASY1 in male meiocytes of the wildtype *Brassica napus* at different stages of prophase I. (B) Co-immunolocalization of SDS and ASY1 in male meiocytes of the *Brassica napus zyp1* mutants at different stages of prophase I. Scale bars=5 μm.

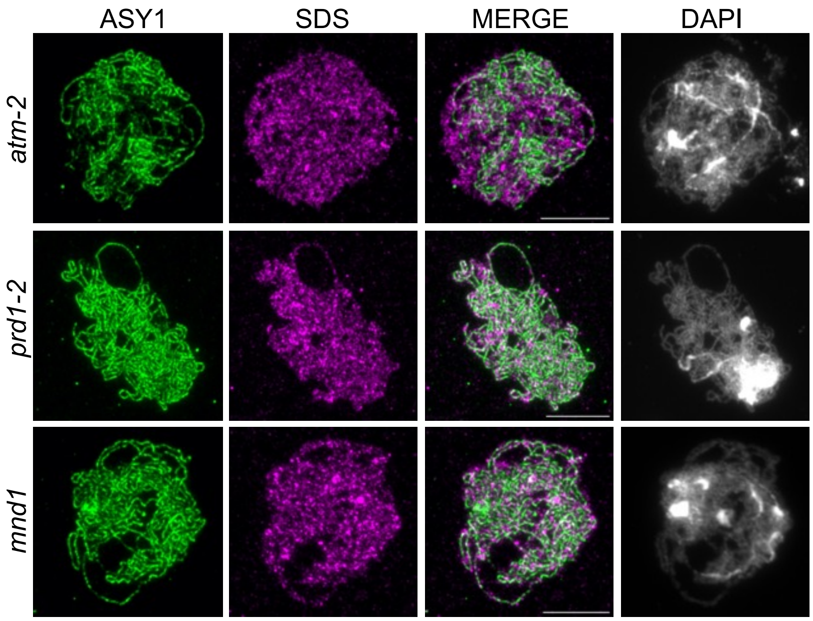


**Supplementary Figure 6. SDS localization on chromosomes is not affected in the absence of ATM, PRD1, or MND1.** Co-immunolocalization of ASY1 and SDS in *atm-2*, *prd1-2*, and *mnd1* mutants. Scale bars=10 µm.

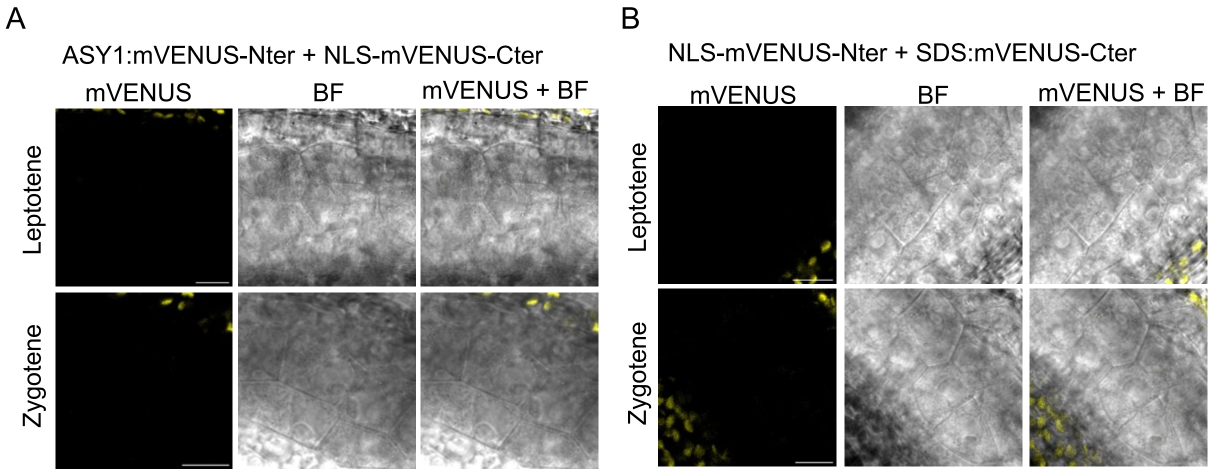


**Supplemental figure 7. Experimental controls for *in planta* BiFC.** (A) The BiFC assay for the combination of ASY1:mVENUS$^{Nter}$ with NLS-mVENUS$^{Cter}$. (B) The BiFC assay for the combination of SDS:mVENUS$^{Cter}$ with NLS-mVENUS$^{Nter}$. All images were captured from Arabidopsis male meiocytes of wildtype during prophase I. Scale bars=10 µm.

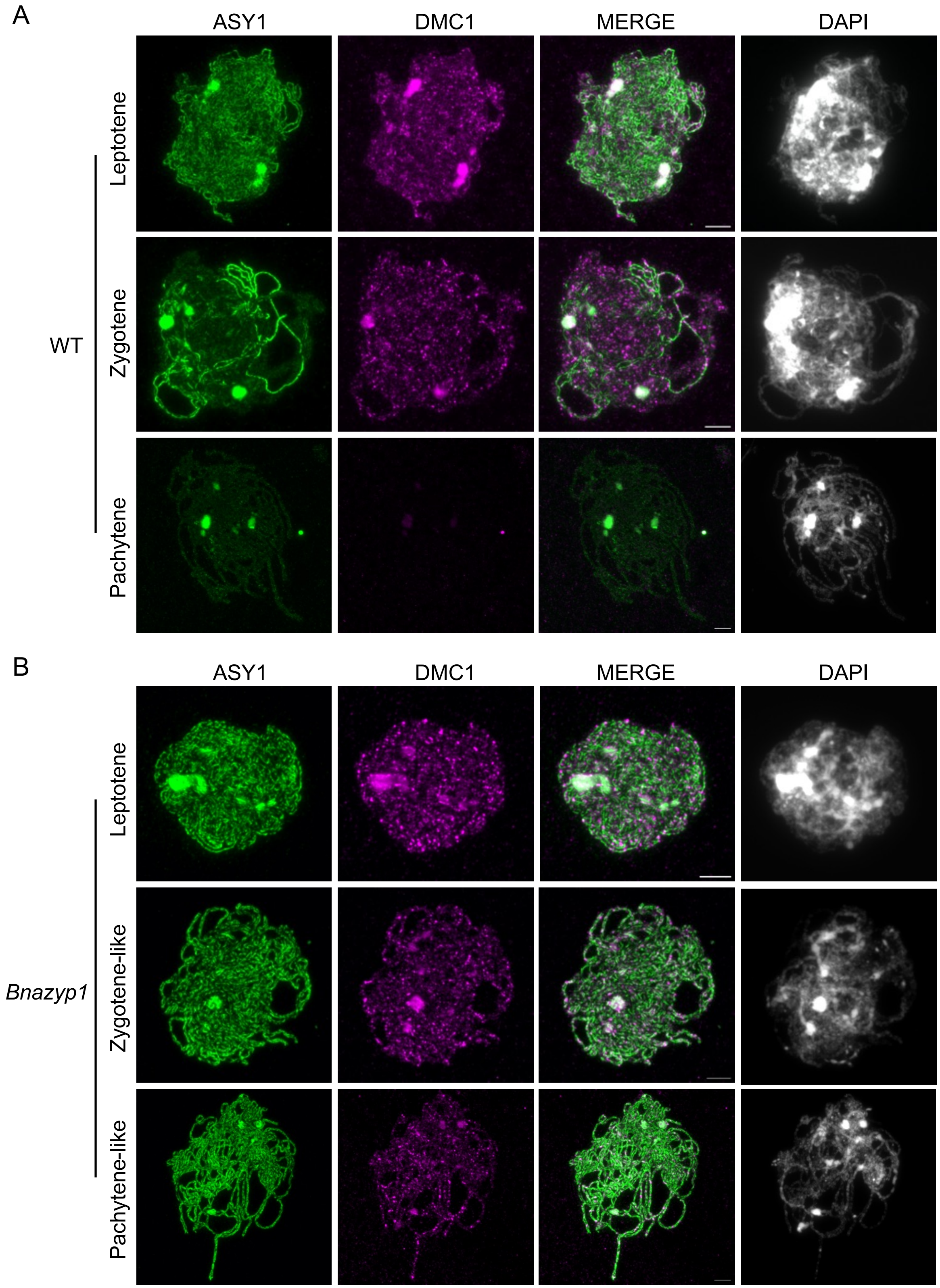


**Supplementary Figure 8. SDS exhibits prolonged stay on meiotic chromosomes in *Brassica napus zyp1* mutants.** (A) Co-immunolocalization of DMC1 and ASY1 in male meiocytes of the wildtype *Brassica napus* at different stages of prophase I. (B) Co-immunolocalization of DMC1 and ASY1 in male meiocytes of the *Brassica napus zyp1* mutants at different stages of prophase I. Scale bars=5 μm.

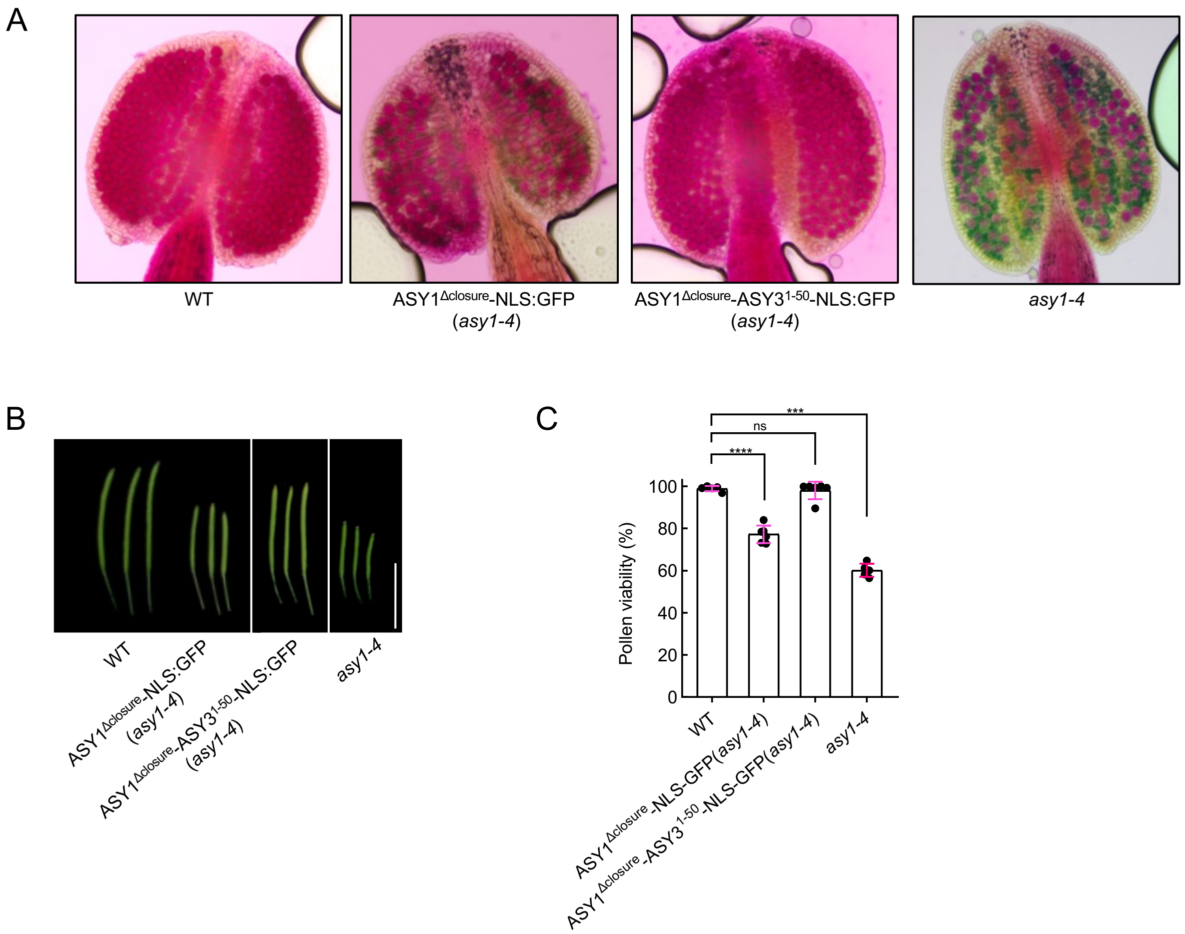


**Supplementary Figure 9. Fertility analysis of *asy1* mutants haboring different ASY1 variants.** (A) Pollen staining of WT, *ASY1$^{\Delta closure}$-NLS:GFP* (*asy1-4), ASY1$^{\Delta closure}$-ASY3$^{1-50}$-NLS:GFP* (*asy1-4*) and *asy1-4* plants. (B) Siliques of WT, *ASY1$^{\Delta closure}$-NLS:GFP* (*asy1-4), ASY1$^{\Delta closure}$-ASY3$^{1-50}$-NLS:GFP* (*asy1-4*) and *asy1-4* plants. Bars: 1 cm. (C) Statistical analysis of pollen viability according to (A). ***, $p<0.001$ (Tukey's multiple comparison test).
